# The membrane-associated ubiquitin ligase MARCHF8 promotes cancer immune evasion by degrading MHC class I proteins

**DOI:** 10.1101/2024.11.29.626106

**Authors:** Mohamed I. Khalil, Jie Wang, Canchai Yang, Lexi Vu, Congcong Yin, Smriti Chadha, Harrison Nabors, Daniel Vocelle, Danielle G. May, Rachel J. Chrisopolus, Li Zhou, Kyle J. Roux, Matthew P. Bernard, Qing-Sheng Mi, Dohun Pyeon

## Abstract

The loss of major histocompatibility complex class I (MHC-I) molecules has been proposed as a mechanism by which cancer cells evade tumor-specific T cells in immune checkpoint inhibitor (ICI)-refractory patients. Nevertheless, the mechanism by which cancer cells downregulate MHC-I is poorly understood. We report here that membrane-associated RING-CH-type finger 8 (MARCHF8), upregulated by human papillomavirus (HPV), ubiquitinates and degrades MHC-I proteins in HPV-positive head and neck cancer (HPV+ HNC). MARCHF8 knockdown restores MHC-I levels on HPV+ HNC cells. We further reveal that *Marchf8* knockout significantly suppresses tumor growth and increases the infiltration of natural killer (NK) and T cells in the tumor microenvironment (TME). Furthermore, *Marchf8* knockout markedly increases crosstalk between the cytotoxic NK cells and CD8^+^ T cells with macrophages and enhances the tumor cell-killing activity of CD8^+^ T cells. CD8^+^ T cell depletion in mice abrogates *Marchf8* knockout-driven tumor suppression and T cell infiltration. Interestingly, *Marchf8* knockout, in combination with anti-PD-1 treatment, synergistically suppresses tumor growth in mice bearing ICI-refractory tumors. Taken together, our finding suggests that MARCHF8 could be a promising target for novel immunotherapy for HPV+ HNC patients.

**One Sentence Summary:** Targeting MARCHF8 restores MHC-I proteins, induces antitumor CD8^+^ T cell activity, and suppresses the growth of ICI-refractory tumors.

## INTRODUCTION

Although immune checkpoint inhibitor (ICI) therapy shows promise in treating various cancers, a majority of cancer patients do not respond ^1–3^. Some solid cancers, including pancreatic, prostate, and head and neck cancers (HNCs), are highly resistant to ICI therapy ^1,4,5^. Human papillomavirus (HPV)-positive HNC (HPV+ HNC) incidence has been dramatically increasing recently despite the effective prophylactic HPV vaccines ^6–8^. It has been found that the levels of viral epitopes in HPV+ HNC patients are associated with a better response to ICI therapy ^9,10^. However, two phase III trials have shown that the response rates to ICI therapies were either similar between HPV+ and HPV-negative (HPV-) HNC patients or even worse in HPV+ HNC patients ^1,11,12^. ICI-refractory patients have limited CD8^+^ T cell infiltration into tumors and low expression of major histocompatibility complex class I (MHC-I) molecules on tumor cells ^13–16^. Many cancers, including HPV+ HNC, show the downregulation of MHC-I and co-stimulatory molecules to avoid eliciting T-cell responses ^17–20^. The levels of MHC-I in HPV+ HNC cells are consistently lower than in normal cells ^21–24^. Nevertheless, the mechanisms of MHC-I downregulation in HPV+ HNC are largely unknown.

Membrane-associated RING-CH-type finger 8 (MARCHF8), an E3 ubiquitin ligase, has initially been identified as a human homolog of viral K3 and K5 ubiquitin ligases encoded by Kaposi’s sarcoma-associated herpesvirus (KSHV) ^25^. Similar to KSHV K3 and K5, MARCHF8 has been found to facilitate the evasion of the host immune responses ^26–28^ by ubiquitinating and degrading several immunoreceptors such as the major histocompatibility complex II (MHC-II) ^29^ and CD86 ^30^ and the cell adhesion molecules CD44 and CD98 ^31^. MARCHF8 expression is frequently upregulated in esophageal, colorectal, hepatic, and gastric cancer and is associated with tumor progression ^32–35^. We have recently shown that MARCHF8 is significantly upregulated in HPV+ HNC cells through the HPV oncoprotein E6-mediated activation of the MYC/MAX transcription factor complex ^36^. Further, MARCHF8 degrades death receptors to inhibit cancer cell apoptosis ^36^ and degrade CUL1 and UBE2L3 to stabilize the HPV oncoprotein E7 ^37^. These results suggest that MARCHF8 is a potent tumor promoter in HPV+ HNC.

Here, we report that MARCHF8 ubiquitinates and degrades MHC-I proteins and that *Marchf8* knockout suppresses in vivo tumor growth by recruiting and activating CD8^+^ T cells in tumor tissues. *Marchf8* knockout, in combination with an anti-PD-1 inhibitor, significantly improves the survival of mice with ICI-refractory HPV+ HNC. These findings provide a novel insight into virus-induced cancer immune evasion and a promising therapeutic target to treat ICI-refractory HPV+ HNC patients.

## RESULTS

### Cell Surface expression of MHC-I on HPV+ HNC cells is downregulated by HPV oncoproteins

To assess MHC-I expression in HNCs, we analyzed the mRNA levels of human leukocyte antigen (HLA)-A, -B, and -C in HPV+ and HPV-HNC patient samples using our previous gene expression profile data ^38^. Our result showed that mRNA expression in HPV+ and HPV-HNC patients is upregulated compared to the normal individuals (**Supplementary Fig. S1A** - **S1C**). Next, to determine MHC-I protein levels in HNCs, we measured protein levels of pan HLA-A/B/C in HPV+ HNC (SCC2, SCC90, and SCC152) and HPV-HNC (SCC1, SCC9, and SCC19) cells compared to normal immortalized keratinocytes (N/Tert-1) cells by western blotting. The results showed that HLA-A/B/C protein levels are significantly lower in HPV+ HNC cells compared to N/Tert-1 and HPV-HNC cells (**Fig. 1A and 1B**). In contrast, MHC-I protein levels are higher in two HPV-HNC cells (SCC1 and SCC9) than in N/Tert-1 cells (**Fig. 1A and 1B**). To examine if the increase in the total MHC-I protein levels reflects MHC-I expression on the cell surface, we determined the levels of HLA-A/B/C proteins on the HPV+ and HPV-HNC cells compared to N/Tert-1 cells using flow cytometry. Consistent with the total protein levels, HLA-A/B/C proteins on all HPV+ HNC cells were significantly decreased compared to N/Tert-1 cells (**Fig. 1C and 1D**). In contrast, HPV-HNC cells, except for SCC19 cells, did not show any decrease of HLA-A/B/C on the cell surface (**Fig. 1C and 1D**). To further determine if the HPV oncoproteins E6 and/or E7 are sufficient to decrease MHC-I protein levels, we measured total and cell surface HLA-A/B/C protein levels in N/Tert-1 cells expressing HPV16 E6 (N/Tert-1 E6), HPV16 E7 (N/Tert-1 E7), or both HPV16 E6 and E7 (N/Tert-1 E6E7) ^37^ using western blotting and flow cytometry, respectively. Interestingly, all N/Tert-1 cells expressing HPV16 E6, E7, or E6E7 significantly decrease MHC-I protein levels in N/Tert-1 cells (**Fig. 1E – 1H**). These results suggest that HPV16 oncoproteins E6 or E7 are sufficient for reducing MHC-I protein levels in normal keratinocytes.

**Fig. 1.**
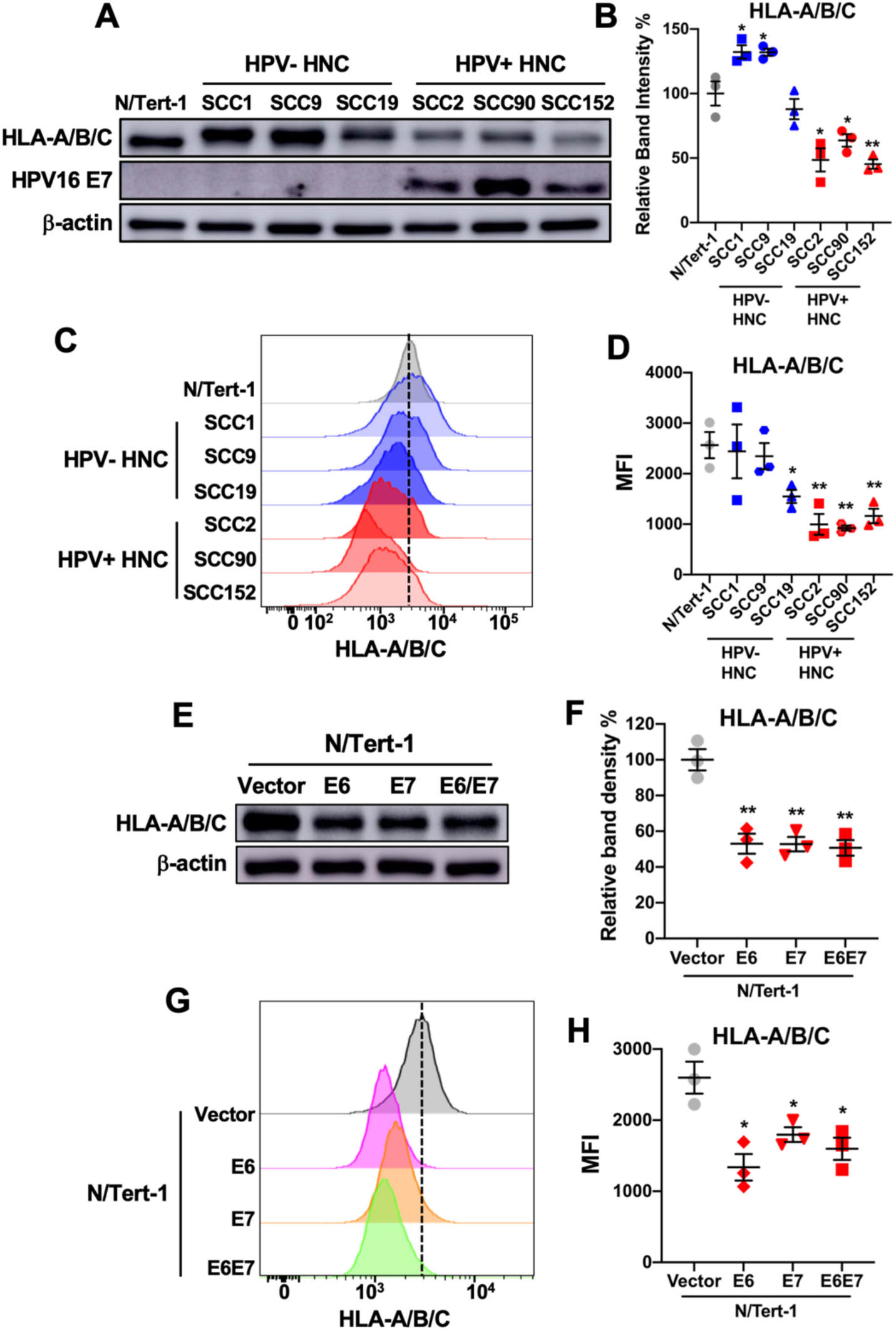
MHC-I expression is downregulated in HPV+ HNC cells. Total protein levels of HLA-A/B/C in HPV-negative (HPV-; SCC1, SCC9, and SCC19) and HPV-positive (HPV+; SCC2, SCC90, and SCC152) HNC cells were determined by western blotting (**A**). Relative band density was quantified using NIH ImageJ (**B**). HPV16 E7 and β-actin were used as viral and cellular controls, respectively. Cell surface expression of HLA-A/B/C (**C** and **D**) on HPV- (SCC1, SCC9, and SCC19) and HPV+ (SCC2, SCC90, and SCC152) HNC cells was analyzed by flow cytometry. Total and cell surface protein levels of HLA-A/B/C in N/Tert-1 cells expressing HPV16 E6, E7, and E6E7 were determined and compared to the vector-only control by western blotting (**E** and **F**) and flow cytometry (**G** and **H**), respectively. Relative band density was quantified using NIH ImageJ (**F**). Mean fluorescence intensities (MFI) of three independent experiments (**D** and **H**) are shown. All experiments were repeated at least three times, and the data shown are means ± SD. *P* values were determined by Student’s *t*-test. \**p* < 0.05, \*\**p* < 0.01.

### MARCHF8 interacts with MHC-I proteins in HPV+ HNC cells

To identify the substrates of the MARCHF8 ubiquitin ligase, we screened MARCHF8 binding proteins in normal immortalized keratinocytes with HPV16 E6 and E7 (N/Tert-1 E6E7) and HPV+ HNC cells (SCC152) using a TurboID-based proximity labeling method ^39^. First, stable cell lines expressing TurboID-MARCHF8 fusion protein or TurboID alone were established and validated by immunofluorescence microscopy (**Fig. 2A**) and western blotting (**Fig. 2B**). Then, biotinylated proteins were pulled down in triplicates and analyzed by mass spectrometry. MARCHF8 interacting proteins were defined with proteins that were pulled down with TurboID-MARCHF8 protein significantly higher (>2-fold, *p* < 0.05) than TurboID alone in both cell lines (**Fig. 2C – 2E, Data file S1**). The quality of our TurboID screen was validated with the identification of those previously known as MARCHF8 binding proteins, such as CDH1, CD44, and TNF-related apoptosis-inducing ligand receptor 1 (TRAIL-R1) (**Fig. 2D and 2E**). Interestingly, two MHC-Iα chains, HLA-A and -C, were identified as MARCHF8 interacting proteins in N/Tert-1 E6E7 and SCC152 cells (**Fig. 2D and 2E**). The Ingenuity Pathway Analysis showed that antigen presentation and ubiquitination are among the top pathways involved in MARCHF8 interactions (**Fig. 2F**). To validate the interaction between MARCHF8 and MHC-I proteins, we pulled down MARCHF8 protein in whole cell lysates from SCC152 cells treated with the proteasome inhibitor MG132 using magnetic beads conjugated with an anti-MARCHF8 antibody. The western blot analyses detected HLA-A/B/C protein in the MARCHF8 protein complex pulled down with an anti-MARCHF8 antibody (**Fig. 2G**). Reciprocally, the co-immunoprecipitation of HLA-A/B/C protein in the same whole cell lysates using an anti-HLA-A/B/C antibody pulled down MARCHF8 protein (**Fig. 2H**), confirming the physical interaction between MARCHF8 and MHC-I proteins. These results suggest that HPV-induced MARCHF8 binds to the MHC-I proteins in HPV+ HNC cells.

**Fig. 2.**
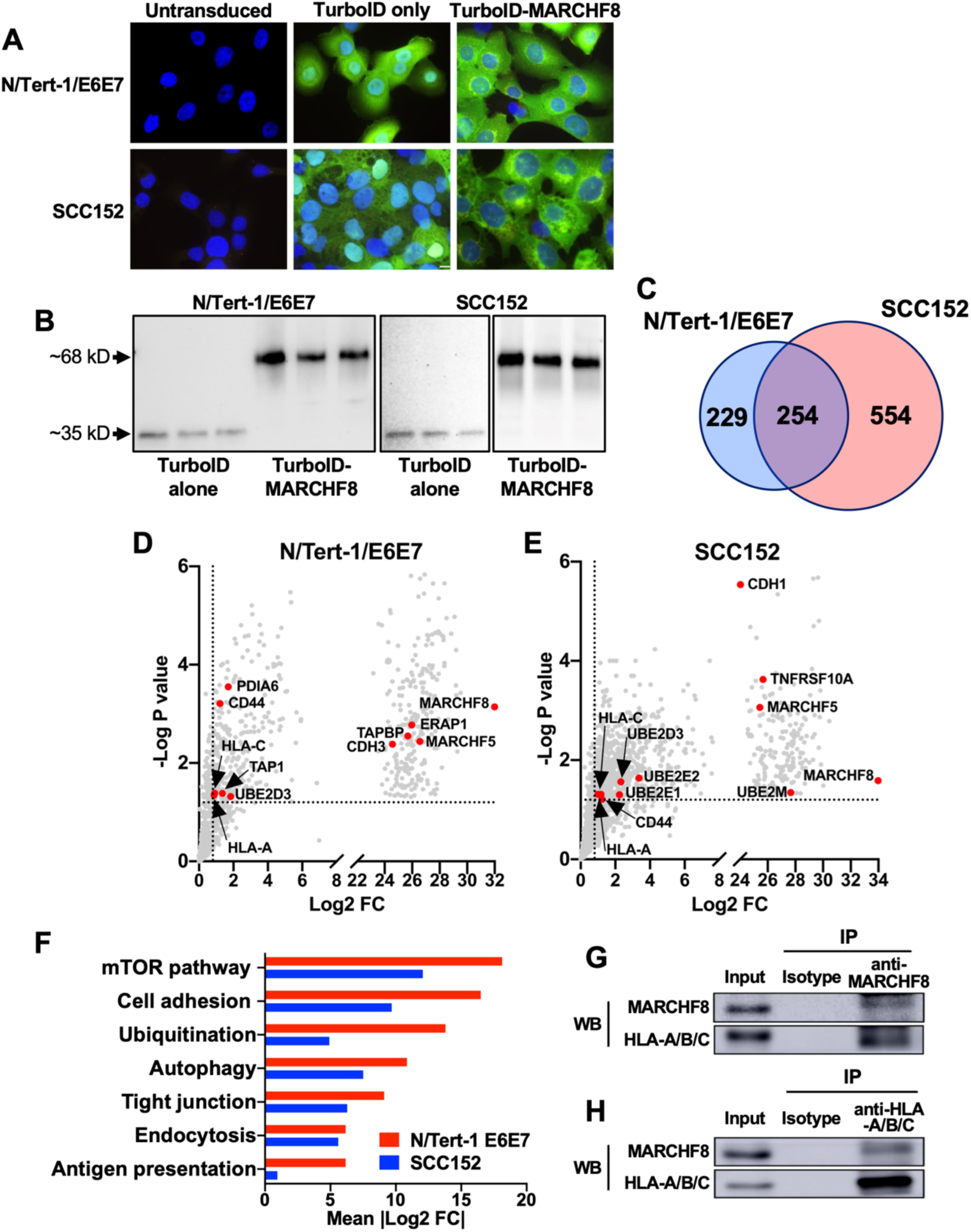
MARCHF8 protein interacts with MHC-I proteins. MARCHF8-TurboID fusion proteins or TurboID proteins alone were stably expressed in N/Tert-1 cells expressing HPV16 oncoproteins E6 and E7 (N/Tert-1/E6E7) and HPV+ HNC cells (SCC152). The TurboID biotinylates the proteins within close proximity. The biotinylated proteins in the cells were separated using streptavidin-coated beads and analyzed using mass spectrometry. Biotinylation by TurboID or TurboID-MARCHF8 was validated by fluorescence microscopy with streptavidin-488 (green) and Hoechst dye (blue) to label DNA (**A**). Expression of TurboID or TurboID-MARCHF8 was validated by western blot with anti-HA (**B**). (**C**) A total of 1,037 proteins enriched with more than a 2-fold increase and *p*-value less than 0.05 with MARCHF8-TurboID compared to TurboID control in both N/Tert-1 E6E7 (**D**) and SCC152 cells (**E**) were defined as potential MARCHF8 binding proteins. The top pathways involved in MARCHF8 interactions were analyzed by Ingenuity Pathway Analysis (**F**). MARCHF8 (**G**) and HLA-A/B/C (**H**) were pulled down from the cell lysate of SCC152 cells treated with MG132 using anti-MARCHF8 (**G**) and anti-HLA-A/B/C (**H**) antibodies, respectively, and analyzed by western blotting. All experiments were repeated at least three times.

### Knockdown of MARCHF8 expression increases MHC-I expression and decreases MHC-I ubiquitination in HPV+ HNC cells

To determine if the upregulated MARCHF8 and its binding to MHC-I are responsible for the decrease of MHC-I protein levels in HPV+ HNC cells, we knocked down MARCHF8 expression in HPV+ HNC cells (SCC152 and SCC2) using multiple shRNAs against MARCHF8 (shR-MARCHF8) delivered by lentiviruses and selected by puromycin treatment. All five SCC152 (**Fig. 3A and 3B**) and three SCC2 (**fig. S2A and S2B**) cell lines with shR-MARCHF8 showed at least a 50% decrease in total MARCHF8 protein levels compared to the control cell lines with scrambled shRNA (shR-scr). Next, we analyzed MHC-I protein levels by western blotting and flow cytometry using an anti-pan HLA-A/B/C antibody. We found that total (**Fig. 3A and 3C, fig. S2A and S2C**) and cell surface (**Fig. 3D and 3E, fig. S2D and S2E**) protein levels of HLA-A/B/C are significantly increased in all SCC152 and SCC2 cell lines by MARCHF8 knockdown. To determine whether the knockdown of MARCHF8 expression affects mRNA expression of MHC-I, we measured mRNA levels of HLA-A, HLA-B, and HLA-C in the SCC152 and SCC2 cell lines by RT-qPCR and found that none of the SCC152 and SCC2 cells show an increase of MHC-I mRNA levels by MARCHF8 knockdown (**fig. S3**). Instead, HLA-A and HLA-B mRNA expression is mostly decreased in SCC152 cells (**fig. S3A – S3C**) but not changed in SCC2 cells by MARCHF8 knockdown (**fig. S3D – S3F**). Next, to determine if MARCHF8 induces ubiquitination of MHC-I proteins, we pulled down ubiquitinated proteins in whole cell lysates from SCC152 cells treated with MG132 using magnetic beads conjugated with an anti-ubiquitin antibody. The results showed that the levels of ubiquitinated HLA-A/B/C proteins were decreased in SCC152 cells by MARCHF8 knockdown, despite the significantly higher levels of total input proteins of HLA-A/B/C, compared to control SCC152 cells with shR-scr (**Fig. 3F**). Our findings suggest that MHC-I protein expression in HPV+ HNC cells is decreased likely by MARCHF8-mediated MHC-I protein ubiquitination.

**Fig. 3.**
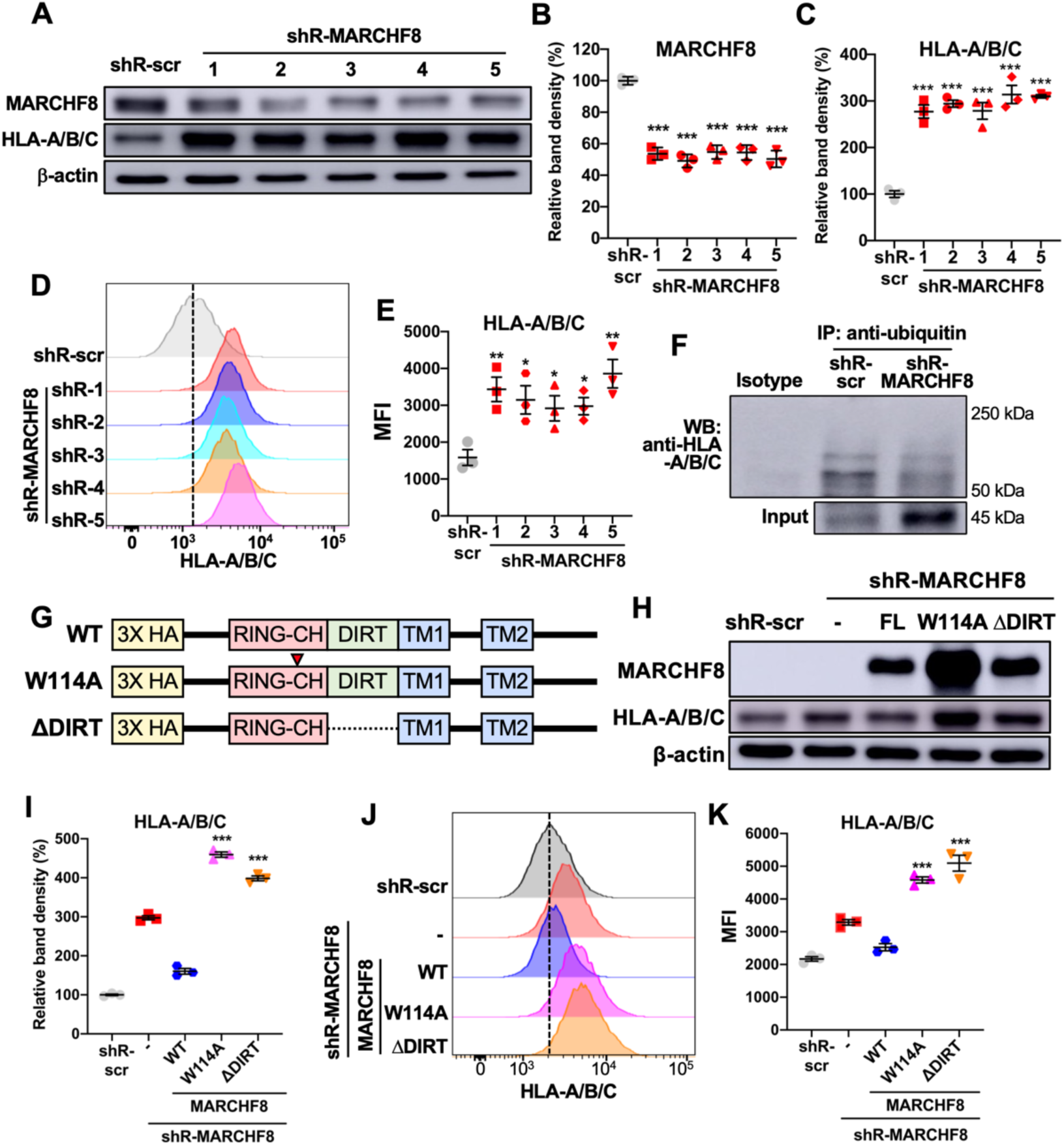
MARCHF8 reduces MHC-I protein levels in HPV+ HNC cells. HPV+ HNC cells (SCC152) were transduced with five lentiviral shRNAs against MARCHF8 (shR-MARCHF8) or scrambled shRNA (shR-scr). MARCHF8 and HLA-A/B/C protein levels were determined by western blotting (**A** - **C**). Relative band density was quantified using NIH ImageJ (**B** and **C**). Cell surface expression of HLA-A/B/C was analyzed by flow cytometry (**D** and **E**). MFI of three independent experiments (**E**) are shown. Ubiquitinated proteins were pulled down from the lysate of SCC152 cells with shR-scr or shR-MARCHF8 (clone 5) using an anti-ubiquitin antibody and analyzed by western blotting (**F**). HA-tagged wild-type (WT) or mutant MARCHF8 constructs were generated with an amino acid substitution in the RING-CH domain (W114A) and deletion of domain in-between RING and transmembrane (ΔDIRT) (**G**). SCC152 cells with shR-MARCHF8 were stably transduced with HA-tagged MARCHF8. MARCHF8 protein expression was determined by western blot using an anti-HA antibody (**H**). Total (**H** and **I**) and cell surface (**J** and **K**) MHC-I levels were measured by western blot and flow cytometry using anti-HLA-A/B/C antibodies. Relative band density was quantified using NIH ImageJ (**I**). The dotted lines are MFI for control (**D** and **J**). The data shown are means ± SD of three independent experiments. *P* values were determined by Student’s *t*-test. \**p* < 0.05, \*\**p* < 0.01, \*\*\**p* < 0.001.

### MHC-I protein degradation by MARCHF8 requires multiple functional domains in MARCHF8 protein

To further elucidate the mechanisms of MHC-I degradation by MARCHF8, we determined the MARCHF8 domains required for MHC-I downregulation by restoring wild-type MARCHF8 or MARCHF8 with mutations in essential domains previously identified ^40,41^. We generated SCC152 cell lines with MARCHF8 knockdown that express HA-tagged wild-type MARCHF8 or MARCHF8 mutants that have an amino acid substitution in the RING-CH domain (W114A) or deletion of domain in-between RING and transmembrane (ΔDIRT) (**Fig. 3G**). We determined MHC-I protein levels in SCC152 cells expressing wild-type or mutant MARCHF8s by western blotting and flow cytometry. Consistent with our results above, the expression of wild-type MARCHF8 significantly decreased the total (**Fig. 3H and 3I**) and cell surface (**Fig. 3J and 3K**) MHC-I protein levels. In contrast, the expression of MARCHF8 mutants, including W114A and ΔDIRT, did not decrease the total (**Fig. 3H and 3I**) and cell surface MHC-I (**Fig. 3J and 3K**) protein levels in SCC152 cells with MARCHF8 knockdown. Interestingly, the expression of MARCHF8 mutants, W114A and ΔDIRT, did not decrease MHC-I expression but rather further increased the total (**Fig. 3H and 3I**) and cell surface HLA-A/B/C (**Fig. 3J and 3K**) protein levels in SCC152 cells with MARCHF8 knockdown, suggesting potential dominant negative functions of these mutant MARCHF8s. These results indicate that MHC-I protein degradation by MARCHF8 requires RING-CH finger and DIRT domains.

### *Marchf8* knockout in mouse HPV+ HNC cells restores MHC-I expression and suppresses tumor growth in vivo

To determine if inhibiting *Marchf8* expression in HPV+ HNC cells suppresses in vivo tumor growth, we used an immunocompetent syngeneic mouse model of mEERL cells expressing HPV16 E6 and E7 along with *Hras*. We first established *Marchf8* knockout mEERL (mEERL/*Marchf8*^-/-^) cell lines using lentiviral transduction of *Cas9* and one of two small guide RNAs (sgRNAs) against *Marchf8* (sgR-*Marchf8*, clones 2 and 3). mEERL cells transduced with *Cas9* and scrambled sgRNA (mEERL/scr) were used as a control. Both mEERL cell lines transduced with sgR-*Marchf8* showed a ∼75% decrease in MARCHF8 protein levels compared to mEERL/scr cells (**Fig. 4A and 4B**). Consistent with the data from human HPV+ HNC cells presented in **Fig. 3**, mEERL/*Marchf8*^-/-^ (sgR-*Marchf8-2* and sgR-*Marchf8-3*) cells showed significantly upregulated H2-Db (MHC-I haplotype in mEERL cell and C57BL/6J mouse) on the cell surface compared to mEERL/scr cells (**Fig. 4C and 4D**). Next, to determine whether *Marchf8* knockout suppresses tumor growth in vivo, either two mEERL/*Marchf8*^-/-^ (sgR-*Marchf8-2* and sgR-*Marchf8-3*) or mEERL/scr cells were subcutaneously injected into syngeneic C57BL/6J mice. Tumor growth was monitored twice a week for 12 weeks. All ten mice injected with mEERL/scr cells showed vigorous tumor growth (**Fig. 4E – 4H**) and succumbed to tumor burden in ∼7 weeks post-injection (**Fig. 4I**). In contrast, the majority of the mice, each injected with either of two mEERL/*Marchf8*^-/-^ (sgR-*Marchf8-2* and sgR-*Marchf8-3*) cells, showed significantly delayed or no tumor growth (**Fig. 4F – 4H**). These results suggest that MARCHF8 is a potent tumor promoter that plays a vital role in HPV+ HNC tumor growth.

**Fig. 4.**
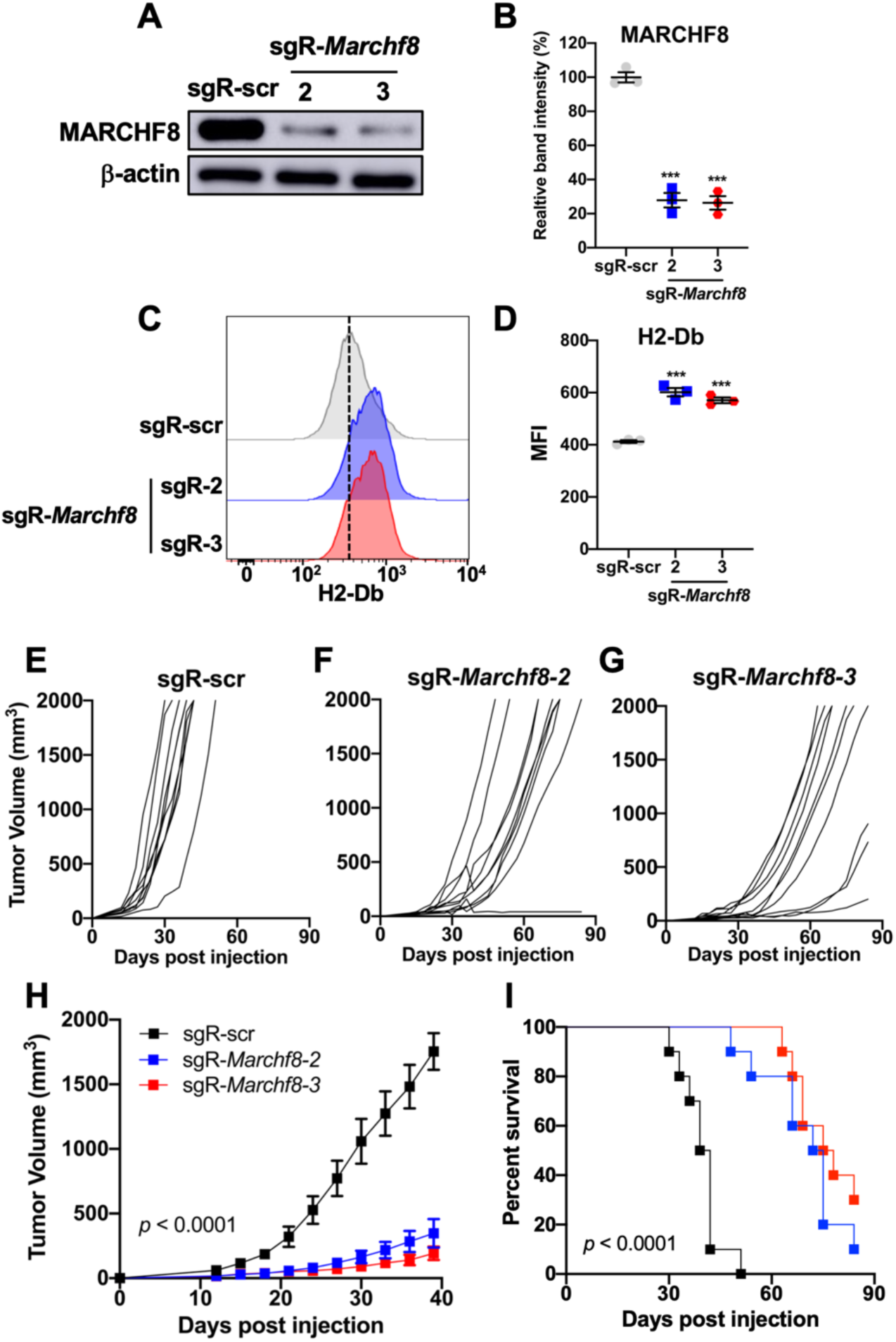
*Marchf8* knockout in HPV+ HNC cells suppresses tumor growth in vivo. mEERL cells were transduced with lentiviral *Cas9* and one of two sgRNAs against *Marchf8* (sgR-*Marchf8-2* and sgR-*Marchf8-3*) or scrambled sgRNA (sgR-scr). MARCHF8 protein levels were determined by western blotting (**A**), and relative band density was quantified using NIH ImageJ (**B**). The data shown are means ± SD of three independent experiments. Cell surface levels of H2-Db protein were measured by flow cytometry (**C** and **D**). The dotted line is MFI for control (**C**). MFI of three independent experiments is shown (**D**). *P* values were determined by Student’s *t*-test. \*\*\**p* < 0.001. mEERL/scr or mEERL/*Marchf8*^-/-^ (sgR-*Marchf8-2* and sgR-*Marchf8-3*) cells were injected into the rear right flank of C57BL/6J mice (*n* = 10 per group). Tumor volume was measured twice a week (**E** - **H**). Survival rates of mice were analyzed using a Kaplan-Meier estimator (**G**). The time to event was determined for each group, with the event defined as a tumor size of 2,000 mm^3^. The data shown are means ± SD. *P* values of mice injected with mEERL/*Marchf8*^-/-^ cells compared with mice injected with mEERL/scr cells were determined for tumor growth (**H**) and survival (**G**) by two-way ANOVA analysis. Shown are representative of two independent experiments.

### *Marchf8* knockout alters immune cell infiltration in the tumor microenvironment (TME) of HPV+ HNC

To characterize immune cell infiltration in the TME altered by *Marchf8* knockout in HPV+ HNC cells, we performed immune cell profiling in tumor tissues harvested from the C57BL/6J mice injected with mEERL/*Marchf8*^-/-^ (sgR-*Marchf8-2* and sgR-*Marchf8-3*) or mEERL/scr cells using single-cell RNA sequencing (scRNA-seq) and flow cytometry. We sorted CD45^+^ cells of the tumor tissues when the average tumor size was 1,000 mm^3^ in the mice. scRNA-seq of 7,965 cells in mEERL/scr tumors, 5,120 cells in mEERL/*Marchf8*^-/-^ tumors (sgR-Marchf8-2), and 6,402 cells in mEERL/*Marchf8*^-/-^ tumors (sgR-Marchf8-3) identified 12 immune cell populations (**Fig. 5A and 5B**). The results showed that macrophages (Clusters #4 and #13), NK/NKT cells (Cluster #10), eosinophils (Cluster #8), and dendritic cell (DC) (Cluster #11) populations were increased in mEERL/*Marchf8*^-/-^ tumors (sgR-*Marchf8-2* and sgR-*Marchf8-3*). In contrast, the B cell (Cluster #5) population was decreased. MDSC (Clusters #1, #2, and #3), CD8 T cells (Cluster #7), and CD4 T cells (Clusters #6 and #9) showed no change or slight increase in mEERL/*Marchf8*^-/-^ tumors (sgR-*Marchf8-2* and sgR-*Marchf8-3*). To validate the key immune cell populations at the cellular levels, we performed a high-dimensional flow cytometry analysis using a 15-antibody panel (**table S1**). Our gating strategy for the immune cell phenotyping is shown in **Fig. S4**. The results showed significant changes in immune cell composition in the TME by *Marchf8* knockout (**Fig. 5C**). Especially, the frequency of CD45^+^ and CD3^+^ cells was dramatically increased in the TME of mEERL/*Marchf8*^-/-^ tumors (sgR-*Marchf8-2* and sgR-*Marchf8-3*) (**Fig. 5D, fig. S5A and S5B**). Although scRNA-seq data showed no change or slight increase in CD4^+^ T cells and CD8^+^ T cells by *Marchf8* knockout, our flow cytometry analysis revealed a significant increase of CD4^+^ T cell and CD8^+^ T cell populations in both mEERL/*Marchf8*^-/-^ tumors, sgR-*Marchf8-2* and sgR-*Marchf8-3* (**Fig. 5C and 5D, fig. S5C**). Additionally, our flow cytometry analysis showed significant increases in NK cell and macrophage infiltration into the TME of mEERL/*Marchf8^-/-^*tumors (sgR-*Marchf8-2* and sgR-*Marchf8-3*) (**Fig. 5D, fig. S5D and S5F**). Furthermore, while scRNA-seq data showed comparable numbers of myeloid-derived suppressor cells (MDSCs) in between mEERL/scr and mEERL/*Marchf8^-/-^* tumors (**Fig. 5B**), we observed that granulocytic MDSC (gMDSC: CD11b^+^ MHCII^-^ CD11c^-^ Ly6G^+^) population was significantly reduced, while monocytic MDSC (mMDSC: CD11b^+^ MHCII^-^ CD11c^-^ Ly6C^+^) population was increased in mEERL/*Marchf8^-/-^* cells (sgR-*Marchf8-2* and sgR-*Marchf8-3*) compared to the sgR-scr tumors (**Fig. 5C, fig. S5E**). These results suggest that the *Marchf8* knockout significantly increases NK cells, T cells, and macrophages potentially remodeling the TME from an immunosuppressive to an immunostimulatory environment.

**Fig. 5.**
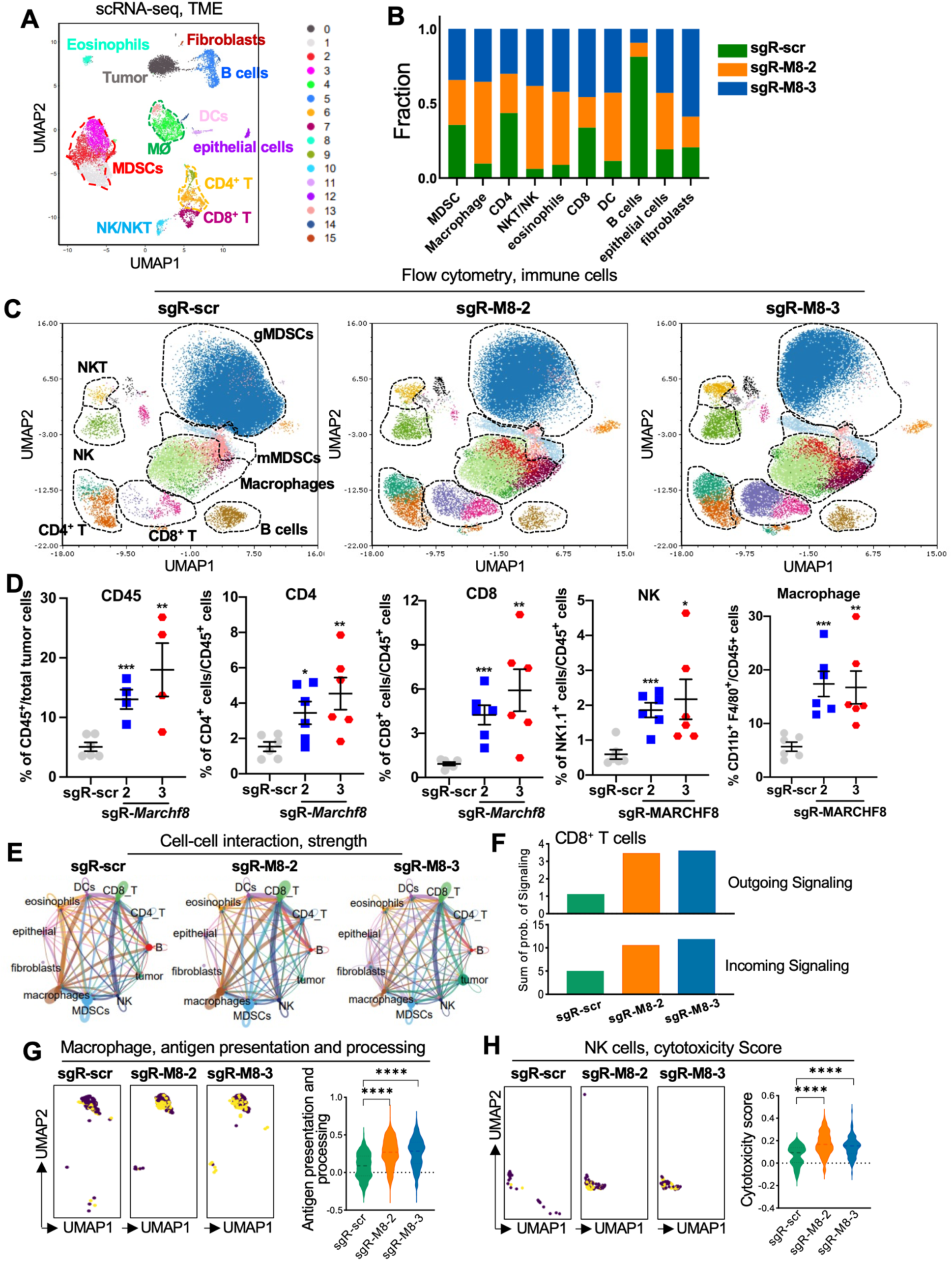
*Marchf8* knockout in HPV+ HNC cells increases NK and CD8^+^ T cells in the TME. Tumors were isolated from C57BL/6J mice injected with mEERL/scr or mEERL/*Marchf8*^-/-^ (sgR-*Marchf8-2* and sgR-*Marchf8-3*) cells. The single cells isolated from the tumor tissues were sequenced for scRNA-seq. The UMAP plots of integrated samples (**A**) and the distribution of each immune cell cluster in *Marchf8* knockout (sgR-Marchf8-2 and sgR-Marchf-3) and scrambled sgRNA (sgR-scr) are shown in the bar graph (**B**). CD45^+^ immune cells were stained with an antibody cocktail and analyzed by flow cytometry. The UMAP plots of flow cytometry data show the color-coded distributions of infiltrating immune cells as indicated (**C**). Dot plots show the frequency of CD45^+^ cells, CD4^+^ and CD8^+^ T cells, NK cells, and macrophages from flow cytometry analysis (**D**). All flow cytometry experiments were repeated at least three times, and the data shown are means ± SD. *P* values were determined by Student’s *t*-test. \**p* < 0.05, \*\**p* < 0.01, \*\*\**p* < 0.001, \*\*\*\**p* < 0.0001. Cell-cell interaction (CCI) was analyzed among different immune cells, and the strength of the CCI was presented as the thickness of the strings (**E**). Bar graphs show the combined outgoing and incoming signaling of CD8^+^ T cells (**F**). UMAP plots show the antigen presentation and processing pathway enrichment in macrophages. The dots in yellow represent the positive in the indicated pathway, and the violin plot shows the score of the antigen presentation and processing in macrophages (**G**). UMAP plots show the cytotoxicity enrichment in NK cells, the dots in yellow represent the positive in the indicated pathway, and the violin plot shows the score of the cytotoxicity enrichment in NK cells (**H**).

To determine how *Marchf8* knockout affects crosstalk among immune cells in the TME, we performed a Cell-Cell Interaction (CCI) analysis using scRNA-seq data. The results showed that the *Marchf8* knockout significantly enhanced the interactions of CD8^+^ T cells with macrophages and NK cells in mEERL/*Marchf8^-/-^* tumors (sgR-*Marchf8-2* and sgR-*Marchf8-3*) compared to mEERL/scr tumors (**Fig. 5E**). Furthermore, both the outgoing signaling from CD8^+^ T cells and incoming signaling to the CD8^+^ T cells was enhanced by *Marchf8* knockout (**Fig. 5F**). These results suggest that *Marchf8* knockout induces active cell-cell communications of CD8^+^ T cells with other immune cells in the TME. To delve deeper into how *Marchf8* affects CD8^+^ T cell interactions with macrophages and NK cells, we performed the KEGG pathway analysis on macrophage and NK cell clusters using scRNA-seq data. Interestingly, macrophage antigen presentation and processing pathways (**Fig. 5G**) and the NK cell cytotoxicity score (**Fig. 5H**) were significantly increased in the mEERL/*Marchf8^-/-^* tumors (sgR-*Marchf8-2* and sgR-*Marchf8-3*). These results suggest that the *Marchf8* knockout may enhance the antigen presentation of macrophages, which is crucial for initiating and sustaining an effective immune response. Additionally, the *Marchf8* knockout enhances the tumor cell-killing activity of NK cells, which is essential in controlling tumor growth and metastasis. Thus, *Marchf8* knockout enhances the crosstalk between CD8^+^ T cells and other immune cells in the TME and may also boost the functional capacities of macrophages and NK cells.

### *Marchf8* knockout in HPV+ HNC cells induces tumor cell-killing functions of CD8^+^ T cells

To determine how *Marchf8* knockout impacts CD8^+^ T cell functions in the TME, we assessed the expression of cytotoxicity-associated genes in CD8^+^ T cells altered by *Marchf8* knockout (**Fig. 6A**). Our results showed a significant increase in the cytotoxicity score of CD8^+^ T cells in the TME of mEERL/*Marchf8^-/-^* tumors (sgR-*Marchf8-2* and sgR-*Marchf8-3*) compared to mEERL/scr tumors (**Fig. 6B**). Specifically, CD8^+^ T cells in the *Marchf8* knockout tumors exhibited significantly increased expression of *Gzmb, Gzma, Prf1, CD44, Ifng,* and *Tbx21* (**Fig. 6C**), indicating that *Marchf8* knockout induces CD8^+^ T cell effector functions for tumor cell-killing. To validate these results, we examined CD8^+^ T cell functions, including proliferation, IFN-γ production, and tumor cell killing. First, to analyze CD8^+^ T cell proliferation, CD8^+^ T cells were isolated from mice injected with mEERL/scr (designated as null CD8^+^ T cells) or mEERL/*Marchf8*^-/-^ cells (designated as primed CD8^+^ T cells). The isolated null or primed CD8^+^ T cells were labeled with carboxyfluorescein diacetate succinimidyl ester (CFSE) and co-cultured with mitomycin C-treated target cells (mEERL/scr or mEERL/*Marchf8*^-/-^ cells). The results showed that the proliferation of CD8^+^ T cells isolated from mice with mEERL/*Marchf8*^-/-^ cells is significantly enhanced compared to CD8^+^ T cells from mice with mEERL/scr cells (**Fig. 6D and 6E**). To determine if *Marchf8* knockout in HPV+ HNC cells activates CD8^+^ T cells, we measured IFN-γ production from CD8^+^ T cells from mEERL/*Marchf8*^-/-^ and mEERL/scr cells using ELISA. The isolated null or primed CD8^+^ T cells were co-cultured with mitomycin C-treated target mEERL/*Marchf8*^-/-^ or mEERL/scr cells. CD8^+^ T cells stimulated with phorbol-12-myristate-13-acetate and ionomycin (PMA/Iono) were used as a positive control. Our results showed that IFN-γ production was significantly increased in primed CD8^+^ T cells co-cultured with mEERL/*Marchf8*^-/-^ cells but not in null CD8^+^ T cells co-cultured with mEERL/scr cells (**Fig. 6F**). Next, to assess the cytolytic capacity of CD8^+^ T cells affected by *Marchf8* knockout in HPV+ HNC cells, we performed a lactate dehydrogenase (LDH) release assay that measures LDH in the cell culture supernatant released by cell lysis. After the isolated null or primed CD8^+^ T cells were incubated with mEERL/scr or mEERL/*Marchf8*^-/-^ cells for 24 hours, the cell culture supernatant was collected, and LDH levels were measured by an enzymatic coupling reaction and absorbance. Interestingly, primed CD8^+^ T cells showed significantly enhanced killing of mEERL/*Marchf8*^-/-^ cells compared to null CD8^+^ T cell killing of mEERL/scr cells (**Fig. 6G**), suggesting that *Marchf8* knockout significantly enhances the antitumor activity of CD8^+^ T cells. In summary, our findings propose that targeting MARCHF8 restores MHC-I expression on HPV+ HNC cells, activates effector CD8^+^ T cells, and suppresses tumor growth (**Fig. 6H**).

**Fig. 6.**
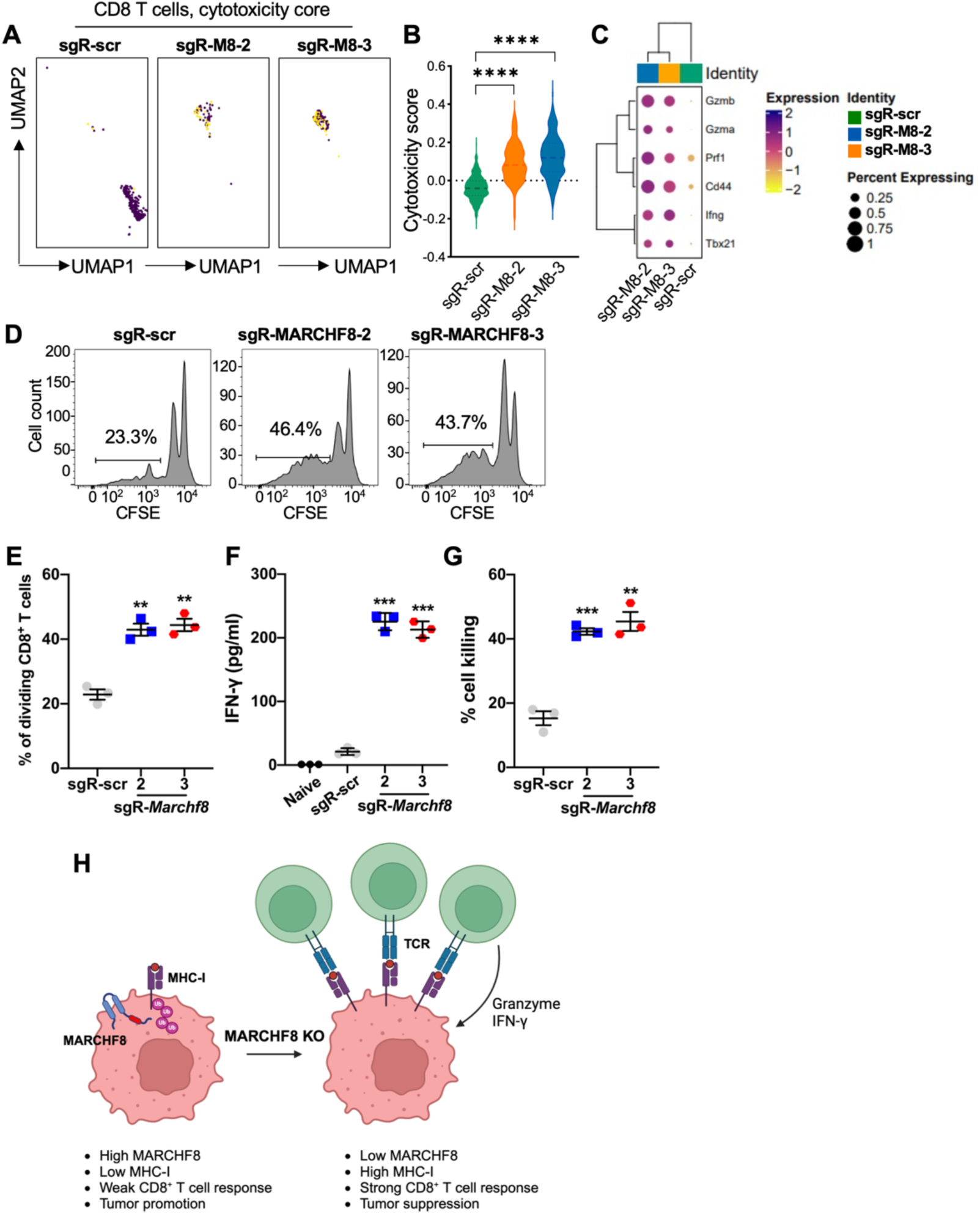
*Marchf8* knockout in HPV+ HNC cells enhances cytotoxic CD8^+^ T cell activation and tumor cell killing. UMAP plots show the cytotoxicity enrichment in CD8^+^ T cells, the dots in yellow represent the positive in the indicated pathway (**A**), and the violin plot shows the score of the cytotoxicity enrichment in CD8^+^ T cells (**B**). The bubble plots show the expression of the indicated genes in CD8^+^ T cells (**C**). Splenic CD8^+^ T cells were isolated from mice injected with mEERL cells using an EasySep mouse CD8^+^ T cells Isolation Kit (StemCell Technologies). The isolated CD8^+^ T cells were labeled with carboxyfluorescein succinimidyl ester (CFSE) and stimulated with anti-CD3/CD28 antibody-coated beads and recombinant mouse IL-2 and co-cultured with either mitomycin-treated mEERL/scr or mEERL/*Marchf8*^-/-^ cells for 3 days. CD8^+^ T cell proliferation was determined by dilution of CFSE intensity using flow cytometry (**D** and **E**). IFN-γ levels in the cell culture supernatant were measured by ELISA (**F**). CD8^+^ T cell cytotoxic activity was determined by the LDH release assay using cell culture supernatant from mEERL/scr or mEERL/*Marchf8*^-/-^ cells co-cultured with CD8^+^ T cells (**G**). All experiments were repeated at least three times, and the data shown are means ± SD. *P* values were determined by Student’s *t*-test. \*\**p* < 0.01, \*\*\**p* < 0.001, ****p < 0.0001. (**H**) Proposed working model illustrating the enhanced tumor cell killing by CD8^+^ T cells following *Marchf8* knockout.

### CD8^+^ T cells are required for tumor suppression by *Marchf8* knockout in HPV+ HNC cells

Given that *Marchf8* knockout upregulates MHC-I in HPV+ HNC cells and activates CD8^+^ T cells for tumor cell killing (**Figs. 4 and 6**), we hypothesized that CD8^+^ T cells are required for tumor suppression by targeting MARCHF8. To test the hypothesis, CD8^+^ T cells were depleted in C57BL/6J mice using an anti-CD8α neutralizing antibody (**Fig. 7A**), as described previously ^21^. Ten doses of an anti-CD8α neutralizing antibody or an IgG2a isotype antibody were intraperitoneally injected into mice with mEERL/*Marchf8*^-/-^ tumors. To validate CD8^+^ T cell depletion in mice, the number of CD8^+^ T cells in splenocytes was counted using an anti-CD8α antibody (clone 2.43) specific to different epitopes from the anti-CD8α neutralizing antibody used in depletion (**table S1**). The results showed that approximately 90% of CD8^+^ T cells were depleted (**Fig. 7B**). Next, we monitored tumor growth and found that CD8^+^ T cells-depleted mice injected with mEERL/*Marchf8*^-/-^ cells showed vigorous tumor growth (**Fig. 7C and 7D**) and succumbed to tumor burden in about 7 weeks post-injection (**Fig. 7E**), despite *Marchf8* knockout. In contrast, *Marchf8* knockout significantly suppresses tumor growth (**Fig. 7C – 7D**) and extended survival (**Fig. 7E**) of the isotype control mice injected with mEERL/*Marchf8*^-/-^ cells compared to the mice injected with mEERL/scr. These results suggest that CD8^+^ T cells are required for tumor suppression by *Marchf8* knockout.

**Fig. 7.**
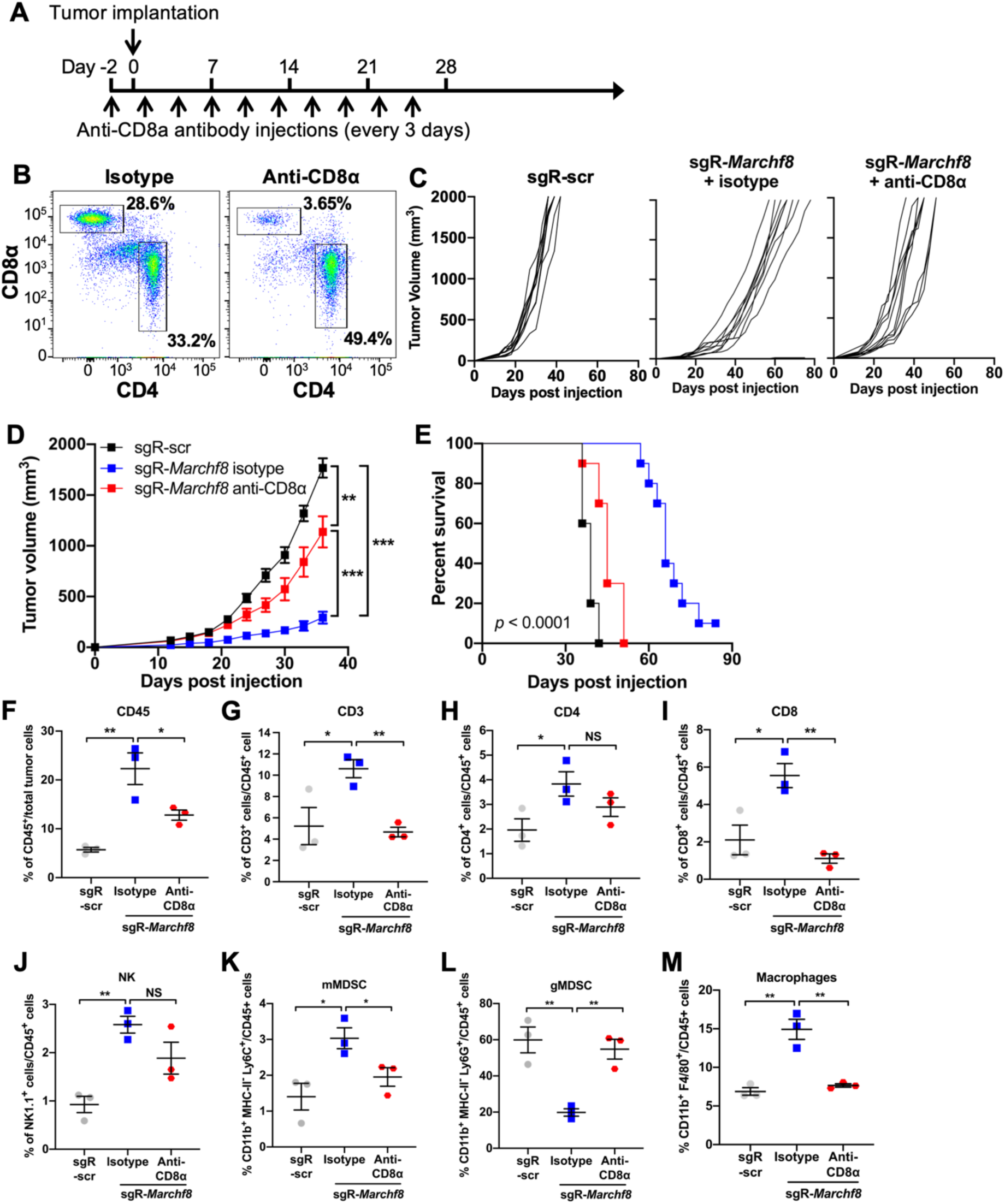
CD8^+^ T cell depletion abrogates tumor suppression by *Marchf8* knockout. C57BL/6J mice were injected with either rat IgG2b (rIgG2b) isotype or anti-mouse CD8α neutralizing (clone 2.43) antibodies. Each mouse received 10 doses of the antibody (100 µg each) starting two days before the injection of mEERL/scr or mEERL/*Marchf8*^-/-^ cells (**A**). CD8^+^ T cell depletion was validated by analyzing CD4^+^ and CD8^+^ T cells in splenocytes (**B**). mEERL/scr or mEERL/*Marchf8*^-/-^ (sgR-*Marchf8* clone2) cells were injected into the rear right flank of C57BL/6J mice (*n* = 10 per group). IgG2b isotype or anti-CD8α antibodies were injected into the mice with mEERL/*Marchf8*^-/-^ cells. Tumor volume was measured twice a week (**C** and **D**). Survival rates were analyzed using a Kaplan-Meier estimator, as described above (**E**). The data shown are means ± SD. *P* values were determined by two-way ANOVA analysis. \*\**p* < 0.01, \*\*\**p* < 0.001. The single cells isolated from tumors were stained with an antibody cocktail and analyzed by flow cytometry. Dot plots show the frequency of the CD45^+^ cells (**F**), CD3^+^ cells (**G**), CD4^+^ (**H**), and CD8^+^ (**I**) T cells, NK cells (**J**), mMDSC (**K**), gMDSC (**L**), and macrophages (**M**). All experiments were repeated at least three times, and the data shown are means ± SD. *P* values were determined by Student’s *t*-test. \**p* < 0.05, \*\**p* < 0.01.

### CD8^+^ T cell depletion restores myeloid cell compositions in the TME altered by *Marchf8* knockout in HPV+ HNC cells

Given that CD8^+^ T cells interact with other immune cells by *Marchf8* knockout (**Fig. 5E**), CD8^+^ T cell depletion may alter other immune cell populations. To test this hypothesis, we assessed immune cells in the TME from CD8^+^ T cell-depleted or control C57BL/6J mice injected with mEERL/*Marchf8^-/-^* cells by flow cytometry as described above. Our UMAP analysis showed that *Marchf8* knockout consistently increases CD45^+^ cells, CD3^+^ cells, CD4^+^ T cells, CD8^+^ T cells, NK cells, and macrophages (**Fig. S6A**). In contrast, CD8^+^ T cell depletion significantly decreased the numbers of CD45^+^ cells (**Fig. 7F, fig. S6B**) and CD3^+^ T cells (**Fig. 7G, fig. S6C**) along with CD8^+^ T cells (**Fig. 7I, fig. S6D**). CD4^+^ T cells (**Fig. 7H, fig. S6D**) and NK cells (**Fig. 7J, fig. S6E**) also decreased, although the decrease was not statistically significant. Interestingly, CD8^+^ T cell depletion in mEERL/*Marchf8*^-/-^ tumors reverses the numbers of MDSCs and macrophages similar to mEERL/scr tumors, increasing the number of gMDSCs (**Fig. 7L, fig. S6F**) and decreasing the number of mMDSCs (**Fig. 7K, fig. S6F**) and macrophages (**Fig. 7M, fig. S6G**). These results suggest that CD8^+^ T cells are critical for converting the immunosuppressive TME to the immunostimulatory TME by *Marchf8* knockout.

### *Marchf8* knockout in HPV+ HNC cells induces CD8^+^ T cell exhaustion, and anti-PD-1 treatment improves the antitumor functions by *Marchf8* knockout

A recent study suggested that intratumoral antigen signaling traps CD8^+^ T cells in the tumor, causing clonal expansion and exhaustion of CD8^+^ T cell populations ^42^. As *Marchf8* knockout in HPV+ HNC cells accumulates CD8^+^ T cells in the TME (**Fig. 5C-D, fig. S5C**), we hypothesize that the CD8^+^ T cells trapped in the *Marchf8* knockout TME undergo exhaustion due to persistent antigen interactions through MHC-I upregulated by *Marchf8* knockout. To test this hypothesis, we examined exhaustion markers, such as PD-1, LAG-3, and TIM-3 on T cells by scRNA-seq data analysis and flow cytometry. The results showed that both mRNA (**Fig. 8A**) and protein (**Fig. 8B – 8D**) levels of PD-1 (*pdcd1*), LAG-3 (*Lag3*), and TIM-3 (*Havcr2*) are highly increased in CD8^+^ T cells by the *Marchf8* knockout. These results suggest that intratumoral antigen signaling activated by *Marchf8* knockout traps CD8^+^ T cells and induces their tumor cell-killing, leading to CD8^+^ T cell exhaustion over time ^42^.

**Fig. 8.**
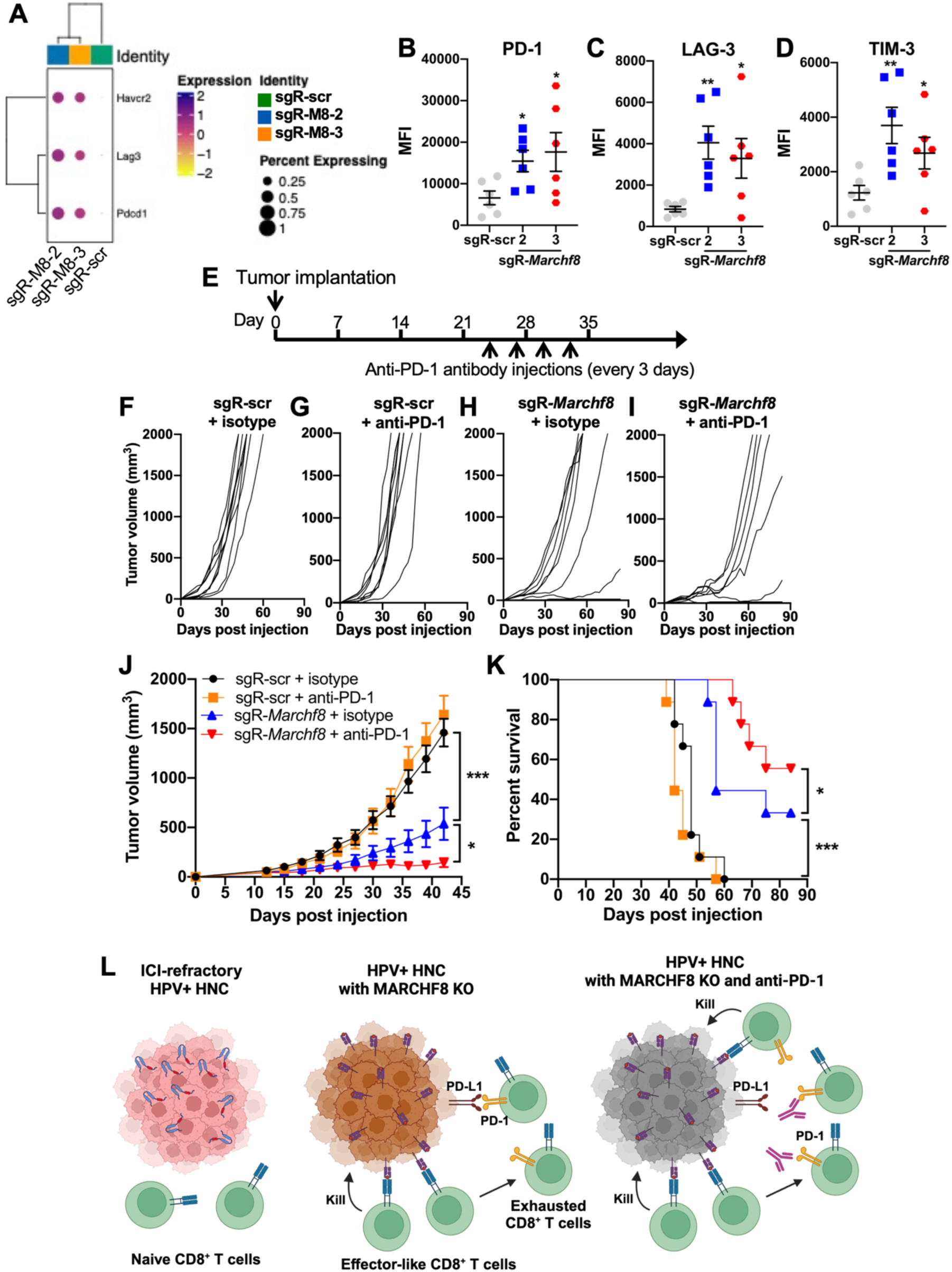
*Marchf8* knockout enhances antitumor activity in combination with anti-PD-1 treatment. The bubble plots show the gene expression changes of CD8^+^ T cells in mEERL/scr or mEERL/*Marchf8*^-/-^ cells, and the genes of T cells exhaustion markers, *Pdcd1*, *Lag3*, and *Havcr2* were labeled (**A**). The protein levels of PD-1 (**B**), LAG-3 (**C**), and TIM-3 (**D**), on CD8^+^ T cells were analyzed by flow cytometry. C57BL/6J mice (*n* = 9 per group) with mEERL/scr or mEERL/*Marchf8*^-/-^ cells were injected with four doses (200 µg each) of either rIgG2a isotype or anti-PD-1 antibodies (clone RMP1-14), starting from day 24 (**E**). Tumor volume was measured twice a week (**F** - **J**). Survival rates were analyzed using a Kaplan-Meier estimator (**K**). The time to event was determined for each group, with the event defined as a tumor size larger than 2,000 mm^3^. The data shown are means ± SD. *P* values were determined by two-way ANOVA for tumor growth (**J**) and survival (**K**). \**p* < 0.05, \*\*\**p* < 0.001. (**L**) The proposed working model illustrates that the *Marchf8* knockout enhances tumor cell killing by CD8^+^ T cells. Additionally, the combination with an immune checkpoint inhibitor (ICI) further improves tumor cell killing by CD8^+^ T cells.

To test if blocking PD-1 rejuvenates the exhausted CD8^+^ T cells and further enhances *Marchf8* knockout-mediated antitumor activity, the mice injected with mEERL/*Marchf8^-/-^* or mEERL/scr cells were treated with four doses of an anti-PD-1 (clone #RMP1-14) or IgG2b isotype antibody every three days starting from 24 days post injection when the average tumor size in mice with mEERL/scr cells was 200 mm^3^ (**Fig. 8E**). Tumor growth was monitored twice a week for 12 weeks. All control mice with mEERL/scr cells showed vigorous tumor growth (**Fig. 8F and 8G**) and succumbed to tumor burden in about eight weeks post-injection (**Fig. 8K**) regardless of the treatment with an anti-PD-1 antibody or an IgG2b isotype antibody. These results suggest that our mEERL mouse model represents the ICI-refractory HPV+ HNCs, showing that ICI treatment did not affect tumor growth and mouse survival. In contrast, the mice with mEERL/*Marchf8*^-/-^ (sgR-*Marchf8-2*) cells consistently showed significant tumor suppression (**Fig. 8H**). Interestingly, the combination of *Marchf8* knockout and anti-PD-1 antibody treatment synergistically suppressed tumor growth (**Fig. 8I and 8J**) and further improved mouse survival, resulting in three out of nine mice with tumor-free (**Fig. 8K**). Notably, tumor growth in four mice with mEERL/*Marchf8*^-/-^ tumors was remitted after the first dose of anti-PD-1 antibody treatment at day 24 (**Fig. 8I**). Our results suggest that anti-PD-1 antibody treatment overcomes CD8^+^ T cell exhaustion induced by *Marchf8* knockout and that MARCHF8 could be an effective target in combination with ICI therapy for treating ICI-refractory HPV+ HNC patients (**Fig. 8L**).

## DISCUSSION

MHC-I molecules are among the most common targets for immune evasion in cancers due to their vital role in antitumor CD8^+^ T cell responses by presenting antigenic peptides ^22,43–48^. Immune evasion through downregulation of MHC-I impairs antitumor immune responses and attenuates immunotherapies that reactivate antitumor CD8^+^ T cells by ICIs ^49,50^. Thus, cancer cells develop various mechanisms to hinder MHC-I antigen presentation, and a better understanding of how cancer cells dysregulate MHC-I antigen presentation is critical.

Previous studies have revealed several mechanisms that downregulate MHC-I expression and prevent antigen presentation in cancer cells. First, genomic alterations by mutation, deletion, and amplification in MHC-I-related genes, including genes for IFN-γ signaling, are common in cancers ^48,51–55^. The loss of heterozygosity (LOH) of at least one of the MHC-I, TAP1, and TAP2 genes in chromosome 6 was found in ∼50% of HNC patients ^56^. PI3K mutations, one of the most common mutations in HNC, downregulates MHC-I by decreasing IFN-γ-induced STAT1 ^57^. These loss-of-function mutations in genes related to MHC-I antigen presentation make cancers refractory to ICIs ^58,59^.

The expression of the MHC-I components is also transcriptionally repressed in several cancers, including prostate, lung, melanoma, and thyroid, by disrupting transcription factors such as NLRC5, IRF2, and NF-κB ^59–62^. Interestingly, while high-risk HPV E7 transcriptionally represses transcription of the MHC-I components, low-risk HPV E7 upregulates their expression ^63^. Epigenetic silencing by DNA methylation and histone modification is another strategy to downregulate MHC-I expression for cancers. The DNA hypermethylation in the MHC-I gene promoters has been found in various cancers, including esophageal squamous cell carcinoma, melanoma, gastric cancer, and breast cancer ^64–67^. Recent studies have shown that the MHC-I components, such as TAP1, TAP2, TAPBP, and B2M, are repressed by polycomb repressive complex 2 (PRC2) ^68,69^. In HNC, reducing H3K27 trimethylation using an EZH2 inhibitor significantly enhances MHC-I antigen presentation ^70^. Thus, inhibitors of epigenetic regulators have been proposed to be used as immunotherapeutics to enhance antitumor immunity ^71,72^. MHC-I expression is also downregulated by microRNAs. Mari *et al.* have shown that increased expression of miR-148a-3p and miR-125-5p in esophageal cancer decreases the mRNA levels of MHC-I and TAP2 ^73^. Similarly, enhanced miR-27a and miR-148a-3p expression reduces MHC-I levels in colorectal cancer by downregulating calreticulin and calnexin, respectively ^74,75^.

MHC-I antigen presentation in cancers is also regulated by posttranslational modification (PTM), which mediates processing antigenic proteins into small peptides and loading them on MHC-I ^76^. On the other hand, PTM also regulates MHC-I trafficking and degradation ^77,78^. MHC-I degradation by ubiquitination is a common immune evasion mechanism, particularly by viruses^79^. MHC-I proteins are ubiquitinated and degraded by human cytomegalovirus US2 and US11 ^80,81^, Epstein Barr virus BDLF3 ^82^, murine γ-herpesvirus 68 K3 ^83^, and KSHV K3 and K5 ^84,85^. KSHV K3 and K5 are membrane-associated E3 ubiquitin ligases that target immune receptors, including MHC-I, for viral immune evasion ^86,87^. While K3 enhances the endosomal degradation of MHC-I via K63 ubiquitination ^88^, K5 inhibits MHC-I phosphorylation ^89^, essential for recycling MHC-I to the cell surface, preventing lysosomal degradation ^90^.

Unlike herpesviruses, HPV does not encode E3 ubiquitin ligases, but its viral oncoproteins E6 and E7 hijack host E3 ubiquitin ligases to target host proteins to facilitate cell cycle progression and prevent apoptosis ^91,92^. We have recently discovered that the HPV oncoproteins upregulate the expression of MARCHF8, a membrane-associated ubiquitin ligase homologous to KSHV K3 and K5, by activating the MYC/MAX transcription factor complex ^25–27,36,93^. MARCHF8, a member of the MARCHF E3 ubiquitin ligase family, is expressed in various tissues, localizes in the plasma and endosomal membranes ^94^, and degrades various membrane proteins ^30,31,95^. We have recently revealed that MARCHF8 ubiquitinates and degrades the death receptors from the TNF receptor superfamily, FAS, TRAIL-R1, and TRAIL-R2 ^36^. Additionally, we report here that MARCHF8 ubiquitinates and degrades MHC-I and that MARCHF8 knockout induces antitumor immune responses by restoring MHC-I expression and activating CD8^+^ T cells in the TME of HPV+ HNC.

FDA-approved ICI therapy has shown promising results in treating patients with previously untreatable cancers ^96,97^. However, clinical trials have shown that most HNC patients do not respond to ICI therapy ^1,98^. Recent studies to better understand the mechanisms of how cancer cells develop resistance to ICIs have revealed diverse strategies to downregulate antigen presentation, increase additional coinhibitory molecules, and create an immunosuppressive TME^99^. While ICI therapy depends on reactivating exhausted T cells, many cancers show limited antigen presentation by downregulating MHC-I and depleting tumor neoantigens to evade T cell recognition ^17,22,99^. Indeed, the non-responders to ICI therapy have low intratumoral CD8^+^ T cell infiltration and frequently have low expression of MHC-I molecules on the tumor cell ^14–16^. The MHC-I levels on HPV+ HNC cells are consistently lower than normal tissue ^21,23,24^. The downregulation of MHC-I is likely one of the key mechanisms of how the response rate of HPV+ HNC patients to ICI therapy is poor despite high PD-L1 expression ^1,11,12^. Thus, restoring MHC-I could be a promising strategy to enhance tumor antigen presentation and antitumor CD8^+^ T cell activity. In this study, targeting MARCHF8 with an anti-PD-1 antibody dramatically suppresses ICI-refractory HPV+ HNC. Taken together, our findings suggest that MARCHF8 could be an effective target for improving ICI therapy for HPV+ HNC patients.

## MATERIALS AND METHODS

### Study design

The main aim of this study is to determine whether the E3 ubiquitin ligase MARCHF8 ubiquitinates and degrades the MHC-I for HPV+ HNC cell immune evasion. We screened the surface and total MHC-I protein expression in HPV+ and HPV-HNC cells using western blotting and flow cytometry. We performed TurboID and immunoprecipitation to determine the interaction of MARCHF8 with MHC-I and the ubiquitination of MHC-I by MARCHF8. We generated human and mouse HPV+ HNC cells with knockdown and knockout of *MARCHF8* to determine the function of MARCHF8 in MHC-I degradation, in vivo tumor growth, and immune cell profiling. Using a combinatory approach of scRNA-seq and high-dimensional flow cytometry, we investigated the role of MARCHF8 in remodeling the immunosuppressive TME to immunostimulatory. The key functions of CD8^+^ T cells in tumor suppression and antitumor immunity were tested by CD8^+^ T cell depletion using a CD8α neutralizing antibody. An anti-PD-1 antibody was used to block PD-1 on CD8^+^ T cells in mice injected with cells with *Marchf8* knockout or scrambled control. The mice were randomly allocated to the experimental groups without blinding procedures. The number of replicates in each experiment is indicated in the figure captions with appropriate statistical analyses.

### Cell lines

HPV+ HNC (SCC2, SCC90, and SCC152) and HPV-HNC (SCC1, SCC9, and SCC19) cells were purchased from the American Type Culture Collection (ATCC) (Manassas, VA), and 293FT cells were purchased from Thermo Fisher (Waltham, MA). These cells were cultured in Dulbecco’s modified Eagle’s medium (DMEM) supplemented with 10% fetal bovine serum (FBS) and penicillin/streptomycin (Thermo Fisher) as described ^100–103^. The N/Tert-1 cell lines were obtained from Dr. Iain Morgan and maintained in keratinocyte serum-free medium supplemented with epidermal growth factor (EGF), bovine pituitary extract, and penicillin/streptomycin as previously described ^104–106^. The mouse oropharyngeal epithelial (MOE) cell line mEERL was obtained from Dr. William Spanos and cultured in E-medium (DMEM and F12 media supplemented with 0.005% hydrocortisone, 0.05% transferrin, 0.05% insulin, 0.0014% triiodothyronine, 0.005% EGF, and 2% FBS) as previously described ^107,108^.

### Flow cytometry

Single cells were isolated from tumor tissue, counted, and analyzed using specific antibodies listed in **table S1**, as previously described ^21^. Cell viability assay was performed by staining cells with Zombie NIR fixable viability dye (BioLegend). Cells for immune cell profiling were stained with antibodies using BD Brilliant staining buffer and analyzed on either a five-laser Cytek Aurora spectral cytometer or a three-laser LSRII flow cytometer (BD Biosciences) in the MSU Flow Cytometry Core Facility. High dimensional reduction and clustering analysis on flow cytometric data was performed using FCS Express (v7) and analyzed utilizing the following pipeline to evaluate global TIL changes in response to treatments for each experimental day separately. First, data was scaled using automatic scaling, then normalized by standard deviation, centered by subtracting the mean, and rescaled to a minimum value of 0 and a maximum value of 1. This ensured a unimodal distribution of the data and that dim populations could be resolved. The data was then gated manually to remove debris, doublets, dead cells, and CD45-negative populations. Next, a UMAP analysis was performed with the following settings to visualize the different sub-populations in distinct groups: all files used, all phenotype parameters used, except for CD45 and Zombie NIR in the FSC files, Neighbors = 50, Minimum Low Dim Distance = 0.9 or 1, Iterations = 500, Random Seed = 886341810. After the UMAP analysis, FlowSOM was used to cluster the data. FlowSOM settings are as follows: all files used, New Scaling = unchecked, Cluster Centroids Initialization Method = Random Cells, Training Decay Function = linear, 2D Grid Neighborhood Function = Boxcar, 2D Grid Distance Metric = Chebyshev, Coarse Training Options = 20, Fine-Tune training Options = 10, Automatic Neighborhood Spread = Checked, Minimum Spanning Tree = Batch SOM Cluster Assignments, Number of Clusters = 18, Sampling Fraction = 1, Number of Samplings = 100, Clustering Algorithms = Hierarchical, Random Seed = 898787008. Next, clusters were assigned a designation based on their phenotype. UMAP scatterplots are shown with color-coded FlowSOM clusters. Clusters were manually annotated based on expression patterns of cellular markers, and dotted lines were drawn to highlight clusters with similar immunophenotypes. Manual gating was performed using FlowJo (v10) to quantify percentages of live cell and immune cell populations.

### Quantitative reverse transcription-PCR (RT-qPCR)

Total RNA was isolated using RNeasy Plus Mini Kit (Qiagen, Germantown, MD). First-strand cDNA was synthesized from 2 µg of total RNA using reverse transcriptase (Roche, Mannheim, Germany). Quantitative PCR (qPCR) was performed in a 20 µl reaction mixture containing 10 µl of SYBR Green Master Mix (Applied Biosystems), 5 µl of 1 µM primers, and 100 ng of cDNA templates using a Bio-Rad CFT Connect thermocycler. Data were normalized to glyceraldehyde 3-phosphate dehydrogenase (GAPDH) or β-actin. Primers used in qPCR (**table S2**) were synthesized by IDT.

### TurboID screen

All TurboID plasmids were constructed using the In-Fusion Cloning system (Takara Bio). HA-tagged TurboID gene (3xHA-TurboID) was amplified from 3xHA-TurboID-NLS pcDNA3 (Addgene, #107171) and inserted into empty pCW57.1 vector (Addgene, #41393) using the NheI and BamHI restriction enzyme (RE) sites, with the addition of an AgeI RE site built into the 3’ primer. pCW57.1-3xHA-TurboID was used as the control plasmid. The human *MARCHF8* gene (Horizon Discovery, MHS6278-202807570) was amplified by PCR and inserted into pCW57.1-3xHA-TurboID at AgeI and BamHI RE sites. SCC152 and N/Tert-1/E6E7 cell lines stably expressing 3xHA-TurboID-MARCHF8 or 3xHA-TurboID alone as control were generated using lentiviral transduction and puromycin selection. 293T cells (ATCC, CRL-3216) were transfected with each construct and third-generation lentiviral packaging plasmids (Cell BioLabs, VPK-206) using Lipofectamine 3000 (Thermo Fisher) according to the manufacturer’s recommendation. The transfected cells were incubated at 37°C for 6 hours, replenished with fresh medium, and further incubated at 37°C for 72 hours. The culture media was filtered through a 0.45-μm filter and added to the cells with polybrene (4 µg/ml; Santa Cruz Biotechnology, Dallas, TX). At 72 hours after transduction, stable cell lines were selected using puromycin (0.5 µg/ml; Thermo Fisher) and verified for TurboID-MARCHF8 fusion protein or TurboID expression and their localization using immunofluorescence and western blotting. For immunofluorescence, cells grown on glass coverslips were fixed in 3% (wt/vol) paraformaldehyde/phosphate-buffered saline (PBS) for 10 min and permeabilized by 0.4% (wt/vol) Triton X-100/PBS for 15 min. For labeling fusion proteins, a mouse anti-hemagglutinin (HA) antibody was used (1:1000; 12CA5; Covance). The primary antibody was detected using Alexa Fluor 488-conjugated or 568–conjugated goat anti-mouse (1:1000; A11001 or A11004; Thermo Fisher Scientific). Alexa Fluor 488–conjugated or 568-conjugated streptavidin (1:1000; S32354 or S11226; Thermo Fisher Scientific) was used to detect biotinylated proteins. DNA was detected with Hoechst dye 33342. Coverslips were mounted using 10% (wt/vol) Mowiol 4-88 (Polysciences). Epifluorescence images were captured using a Nikon Eclipse NiE (20×/0.75 Plan Apo Nikon objective) microscope. To analyze total cell lysates by western blot, 1.2 × 10^6^ cells were lysed in SDS–PAGE sample buffer, boiled for 5 min, and sonicated to shear DNA. Proteins were separated on 4–20% gradient gels (Mini-PROTEAN TGX; Bio-Rad, Hercules, CA) and transferred to nitrocellulose membrane (Bio-Rad). After blocking with 10% (vol/vol) adult bovine serum and 0.2% Triton X-100 in PBS for 30 min, the membrane was incubated with rabbit anti-HA antibody (1:20000; ab9110; Abcam) overnight, washed with PBS and detected using horseradish peroxidase (HRP)–conjugated anti-rabbit (1:40,000; G21234; Invitrogen). The signals from antibodies were detected using enhanced chemiluminescence via a Bio-Rad ChemiDoc MP System (Bio-Rad, Hercules, CA). Following the detection of HA, the membrane was quenched with 30% H_2_O_2_ for 30 min. To detect biotinylated proteins, the membrane was incubated with HRP-conjugated streptavidin (1:40,000; ab7403; Abcam) in 0.4% Triton X-100 in PBS for 45 min. The stable cell lines were maintained in DMEM (Corning, 10-013-CV) supplemented with 10% FBS. Large-scale TurboID pulldowns were performed in triplicate as described ^109^. Briefly, four 10 cm dishes at 80% confluency were incubated with doxycycline (1 µg/ml) for 18 hours before biotinylation with 50 µM biotin for 4 hours. Cells were rinsed twice with PBS and lysed in 50 mM Tris pH7.4 buffer containing 8 M urea and protease inhibitors (Thermo Fisher, 87785) and DTT, incubated with universal nuclease (Thermo Fisher, 88700), and sonicated to shear DNA further. Lysates were precleared with Gelatin Sepharose 4B beads (GE Healthcare, 17095601) for 2 hours and then incubated with Streptavidin Sepharose High-Performance beads (GE Healthcare, 17511301) for 4 hours. Streptavidin beads were washed four times with 50 mM Tris pH7.4 wash buffer with 8 M urea and resuspended in 50 mM ammonium bicarbonate with 1 mM biotin.

### Mass spectrometry

Protein samples were reduced, alkylated, and digested using filter-aided sample preparation with porcine trypsin (Promega) ^110^. Tryptic peptides were separated by reverse phase XSelect CSH C18 2.5 µm resin (Waters) on an in-line 150 x 0.075 mm column using an UltiMate 3000 RSLCnano system (Thermo Fisher). Peptides were eluted using a 60-min gradient from 98:2 to 65:35 buffer A:B ratio (Buffer A = 0.1% formic acid, 0.5% acetonitrile, Buffer B = 0.1% formic acid, 99.9% acetonitrile). Eluted peptides were ionized by electrospray (2.4 kV), followed by mass spectrometric (MS) analysis on an Orbitrap Fusion Tribrid mass spectrometer (Thermo Fisher). MS data were acquired using the Fourier Transform Mass Spectrometer in profile mode at a resolution of 240,000 over 375 to 1500 m/z. Following higher-energy collisional dissociation activation, MS/MS data were acquired using the ion trap analyzer in centroid mode and normal mass range with a normalized collision energy of 28-31%, depending on the charge state and precursor selection range. Proteins were identified by database search using MaxQuant (Max Planck Institute) label-free quantification with a parent ion tolerance of 2.5 ppm and a fragment ion tolerance of 0.5 Da. Scaffold Q+S (Proteome Software) was used to verify MS/MS-based peptide and protein identifications. Protein identifications were accepted if they could be established with less than 1.0% false discovery and contained at least two identified peptides. Protein probabilities were assigned by the Protein Prophet algorithm ^111^. Protein interaction candidates were considered if identified in at least two replicate runs and enriched over the TurboID-only control.

### Immunoprecipitation (IP) and western blotting

Whole cell lysates were prepared in radioimmunoprecipitation assay buffer (RIPA) buffer (Abcam, Waltham, MA) containing a protease inhibitor cocktail according to the manufacturer’s instructions (Roche, Mannheim, Germany). Total protein concentrations were determined using the Pierce BCA Protein Assay Kit (Thermo Fisher). IP was performed using the Pierce Classic Magnetic IP/Co-IP Kit (Thermo Fisher). Briefly, 25 µl of protein A/G magnetic beads were incubated with 5 µg of specific antibodies (**table S1**) for 2 hours. One mg of the whole cell lysates was incubated with antibody-coupled beads at 4°C overnight. Western blotting was performed with 10 to 20 µg of total protein using antibodies listed in **table S1** as previously described ^112^. Band densities were determined using NIH ImageJ software and normalized to the β-actin band intensity.

### Lentivirus production and transduction

The 3XHA-hMARCHF8-W114A coding sequence was amplified from pCAGGS-3xHA-MARCHF8-W114A (provided by Dr. Yong-Hui Zheng), and the HA-tagged wild-type MARCHF8 coding sequence was generated from pCAGGS-3XHA-hMARCHF8 (provided by Dr. Yong-Hui Zheng) ^41^ using the primers listed in **table S2** and cloned into the SpeI and AgeI RE sites in pLenti6/V5-D-TOPO vector (Addgene, #22945). The DIRT domain was deleted from pLenti6-3xHA-MARCHF8 using the QuikChange II XL Site-Directed Mutagenesis Kit (Agilent, 200522) and the primers listed in **table S2**. The shRNAs targeting human *MARCHF8* were purchased from Sigma-Aldrich. The sgRNAs targeting the mouse *Marchf8* were designed by the web-based software ChopChop ^113^. sgRNAs were synthesized and cloned into the lentiCRISPR v2-blast plasmid (Addgene, #83480) using ligating duplex oligonucleotides containing BsmBI RE sites (IDT). All shRNA and sgRNA sequences are listed in **tables S3 and S4**, respectively. Lentiviruses containing shRNA or sgRNA were produced using 293FT cells with packaging constructs pCMV-VSV-G (Addgene, #8454) and pCMV-Delta R8.2 (Addgene, #12263). The lentiviruses were collected 48 hours post-transfection and concentrated by ultracentrifugation at 25,000 rpm for 2 hours. Cells were incubated with lentiviruses for 48 hours in polybrene (8 µg/ml) and selected with blasticidin (8 µg/ml).

### Mice and tumor growth

C57BL/6J mice were purchased from Jackson Laboratory (Bar Harbor, ME) and maintained following the USDA guidelines. Six to eight-week-old mice were injected with 5 X 10^5^ mEERL cells subcutaneously into the rear right flank (*n* = 10 per group). Tumor volume was measured twice weekly and calculated using the equation: volume = (width^2^ X length)/2. Animals were euthanized when tumor volume reached 2,000 mm^3^, as previously described in ^114^. Conversely, mice were considered tumor-free when no measurable tumor was detected for 12 weeks. Survival curves were generated by Kaplan–Meier analysis standardizing for a tumor volume of 2,000 mm^3^. The Michigan State University Institutional Animal Care and Use Committee (IACUC) approved experiments using live animals according to National Institutes of Health guidelines. CD8^+^ T cells in C57BL/6J mice were depleted by intraperitoneal injection of 100 µg of an anti-CD8α neutralizing antibody (BioXcell, clone 2.43) every three days for 30 days starting two days before tumor cell injection. An IgG2a isotype antibody was used as an isotype control. For anti-PD-1 treatment, C57BL/6J mice were intraperitoneally injected with 200 µg of an anti-PD-1 antibody (BioXcell, clone RMP1-14) every three days for four times starting from 24 days post-injection when the average tumor size in the mice injected with mEERL/scr cells reached 200 mm^3^. An IgG2b isotype antibody was used as an isotype control.

### Intratumoral immune cell isolation

The immune cells were isolated from tumor tissues as previously described ^115^. Tumors were surgically removed from mice, minced into RPMI medium, and incubated with 0.5 mg/ml collagenase IV (Millipore-Sigma) and 1,000 IU/ml of DNaseI (Millipore-Sigma) at 37°C for one hour with vortexing every 15 min. After digestion, cells were resuspended in RPMI supplemented with 10% FBS and filtered through a 40 µM cell strainer. Red blood cells (RBCs) were lysed by adding 1.5 ml of RBC lysis buffer (Millipore-Sigma) for less than 5 min.

### Single-cell RNA sequencing (scRNA-seq)

ScRNA-seq libraries were generated using the 10X Genomics Chromium Single Cell 3’ Reagent Kit (v2 Chemistry) and Chromium Single Cell Controller as described in the previously published study ^116^. Briefly, FACS-sorted cells were loaded into each reaction for gel bead-in-emulsion (GEM) generation and cell barcoding. Reverse transcription of the GEM (GEM-RT) was performed using a Veriti 96-Well Fast Thermal Cycler (Applied Biosystems) at 53 C for 45 min, 85 C for 5 min, and 4°C hold. cDNA amplification was performed after GEM-RT cleanup with Dynabeads MyOne Silane (ThermoFisher Scientific) using the same thermocycler (98 C for 3 min, 98 C for 15 seconds, 67 C for 20 seconds, 72 C for 1 min, for 12 cycle repeats followed by a 72 C for 1 min and a 4 C hold). The amplified cDNA was cleaned up with SPRIselect Reagent Kit (Beckman Coulter, Brea, CA), followed by a library construction procedure, including fragmentation, end repair, adaptor ligation, and library amplification. An Agilent 2100 Bioanalyzer (Agilent, Santa Clara, CA) was used for library quality control. Libraries were sequenced on an Illumina HiSeq4000 using a paired-end flow cell: Read 1, 26 cycles; i7 index, 8 cycles; Read 2, 98 cycles.

### ScRNA-seq data analysis

As has been described in our previous study ^116^, sequenced reads from scRNA-seq libraries were demultiplexed, aligned to the mm10 mouse reference, barcode processed, and Unique Molecular Identifier (UMI) counted using 10X Genomics Cell Ranger (v2.0.1) pipeline. Datasets were subsequently analyzed using the R Seurat package. Principal Component Analysis (PCA) was employed to analyze combined samples. Cells with unique feature counts over 2,500 or less than 200 and/or high mitochondrial genetic content (>5%) were filtered out. A global-scaling normalization method, “LogNormalize” in Seurat, was employed to normalize gene expression. Highly variable genes in each data analysis were identified, and the intersecting top 2,000 genes in each dataset were used for clustering and subsequent downstream analyses. The first 30 principal components (PCs) used the Uniform manifold approximation and projection (UMAP) method to visualize the resulting clusters. The FindAllMarkers function in Seurat was then implemented to identify DEGs between clusters with a Bonferroni adjustment of *P* value <0.05 as a statistical significance threshold.

### T cells isolation and stimulation

Mouse spleens were mechanically disrupted over a 100-µm filter and subjected to magnetic bead negative enrichment for CD8**^+^** T cells using a CD8**^+^** T-cell Isolation Kit (STEMCELL Technologies). Isolated CD8**^+^** T cells were cultured in RPMI (Gibco) containing 10% FBS, L-glutamine, penicillin/streptomycin, and β-mercaptoethanol (50 µM). Isolated CD8**^+^** T cells (1 × 10^6^ cells/ml) were stimulated using Dynabeads Mouse T-Activator CD3/CD28 (Invitrogen). Stimulated cells were incubated in media supplemented with 10 ng/mL of recombinant mouse IL-2 (rmIL-2) (ThermoFisher).

### CD8^+^ T cell proliferation assay

CD8**^+^** T cells were isolated from the mouse spleen and injected with mEERL cells with scramble or *Marchf8* sgRNA using the CD8**^+^** T-cell Isolation Kit (STEMCELL Technologies). The cells were labeled with 3 µM CFSE (Invitrogen). The labeled CD8**^+^** T cells were co-cultured with mitomycin-C treated mEERL cells for 3 days with scramble or *Marchf8* sgRNA in RPMI media containing 10 ng/mL of rmIL-2 and Dynabeads Mouse T-Activator CD3/CD28 for T-cell Expansion and Activation Kit. CD8**^+^** T cells were collected, and dye concentrations were measured using flow cytometry.

### Enzyme-linked immunosorbent assay (ELISA)

Isolated CD8**^+^** T-cells were co-cultured with mitomycin-C-treated mEERL cells with scramble or *Marchf8* sgRNA in RPMI containing 10 ng/mL rmIL-2 and Dynabeads Mouse T-Activator CD3/CD28 for T-cell Expansion and Activation kit (Invitrogen) for 3 days. Cell culture supernatants were collected and used to quantify IFN-γ production by ELISA (Biolegend, #430801) according to the manufacturer’s instructions.

### Lactate dehydrogenase release assay

To measure the CD8**^+^** T-cell cytotoxicity, one million isolated CD8**^+^** T cells (effector) were incubated with 10,000 mEERL cells (target) with scramble or *Marchf8* sgRNA for 24 hours. Cell culture supernatant was collected, and cell cytotoxicity was determined by measuring the lactate dehydrogenase (LDH) concentrations in the cell culture supernatant using the Cytotoxicity Detection Kit (Millipore Sigma). As controls, spontaneous and maximum cell lysis were assessed in the absence of CD8**^+^** T cells and by treating the mEERL cells with 1% SDS, respectively.

### Statistical analysis

Data were analyzed using GraphPad Prism (San Diego, CA) and were presented as mean ± standard deviation. Statistical significance was determined using an unpaired Student’s *t-*test. *P* values <0.05 are considered statistically significant. Distributions of time-to-event outcomes (e.g., survival time) were summarized with Kaplan–Meier curves and compared across groups using the log-rank test with α = 0.01.

## Supplementary Materials

Figs. S1 to S6 for multiple supplementary figures.

Tables S1 to S4 for multiple supplementary tables.

Data file S1 to S3 for multiple data files.

## Acknowledgment

We thank Yong-Hui Zheng, Iain Morgan, John H. Lee, and William C. Spanos for the valuable reagents, and Bardees Foda and members of the Pyeon laboratory for their valuable comments and suggestions.

## Funding

National Institutes of Health grant R01 DE029524 (D.P.)

Michigan State University Global Impact Initiative (D.P.)

MSU Foundation Strategic Partnership Grant (D.P.).

National Institutes of Health grant P20 GM103620 (K.J.R)

National Institutes of Health grant R24 GM137786 (K.J.R)

National Institutes of Health grant P30 GM145398 (K.J.R).

## Author contributions

Conceptualization: M.I.K. and D.P.

Methodology: M.I.K, J.W., D.V., D.G.M., R.J.C., L.Z., K.J.R., M.P.B., and Q.S.M.

Investigation: M.I.K, J.W., C.Y., L.V., C.Y., S.C., H.N., Q.S.M., and D.P.

Visualization: M.I.K, J.W., and D.P.

Project administration: M.I.K. and D.P.

Writing original draft: M.I.K. and D.P.

Writing—review and editing: M.I.K., J.W., C.Y., L.V., K.J.R., M.P.B, Q.S.M., and D.P.

## Competing interests

The authors declare that they have no competing interests.

## Notes

### Competing Interest Statement

The authors have declared no competing interest.

