## Supplemental Information for "The membrane-associated ubiquitin ligase MARCHF8 promotes cancer immune evasion by degrading MHC class I proteins"

Mohamed I Khalil et al.

**The PDF file includes:**

Figs. S1 to S6
Tables S1 to S4

**Other Supplementary Material for this manuscript includes the following:**

Data files S1 to S3

Khalil et al., Fig.S1.

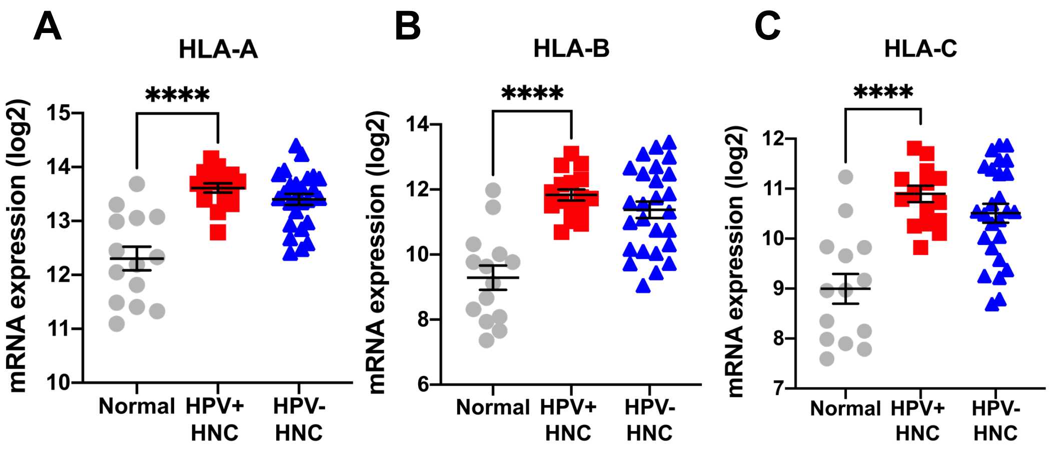


**Fig. S1. mRNA expression levels of HLA-A, HLA-B, and HLA-C in HPV+ and HPV- HNC patient samples.** The HLA-A, HLA-B, and HLA-C mRNA expression levels in micro-dissected human tissue samples from HPV+ (n = 16), HPV- (n = 26) HNC patients, and normal individuals (n = 12) were analyzed using our previous gene expression data (GSE6791) [38] and shown as fluorescence intensity (Log2) (**A-C**). The data shown are means ± SD. *P* values were determined by Student's *t*-test. *****p* < 0.0001.

Khalil et al., Fig. S2.

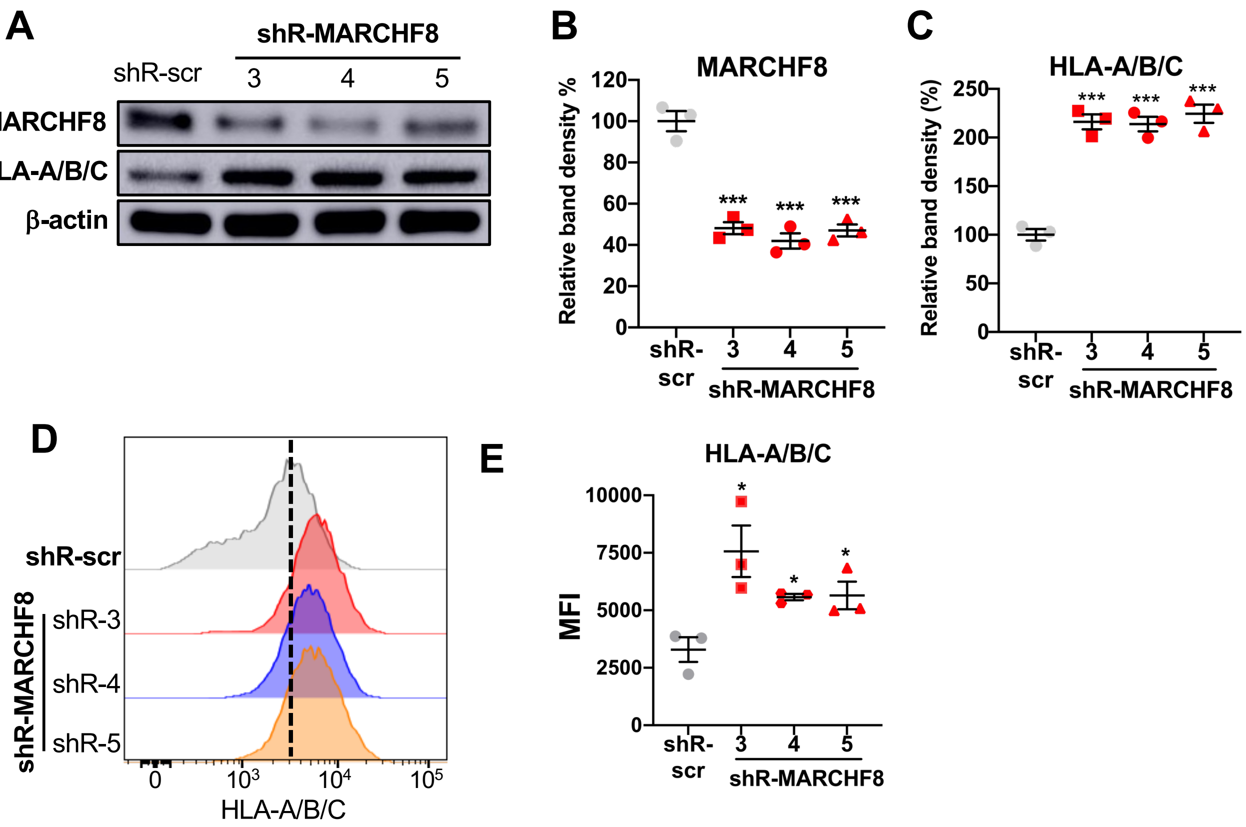


**Fig. S2. MARCHF8 knockdown increases MHC-I protein levels in HPV+ HNC cells.** HPV+ HNC cells (SCC2) were transduced with three lentiviral shRNAs against MARCHF8 (shR-MARCHF8) or scrambled shRNA (shR-scr). Protein levels of MARCHF8 and HLA-A/B/C were determined by western blotting (**A** - **C**). Relative band density was quantified using NIH ImageJ (**B** and **C**). Cell surface expression of HLA-A/B/C (**D** and **E**) proteins was analyzed by flow cytometry. MFI of three independent experiments is shown (**E**). The data shown are means ± SD of three independent experiments. *P* values were determined by Student's *t*-test. $^*p < 0.05$, $^{***}p < 0.001$.

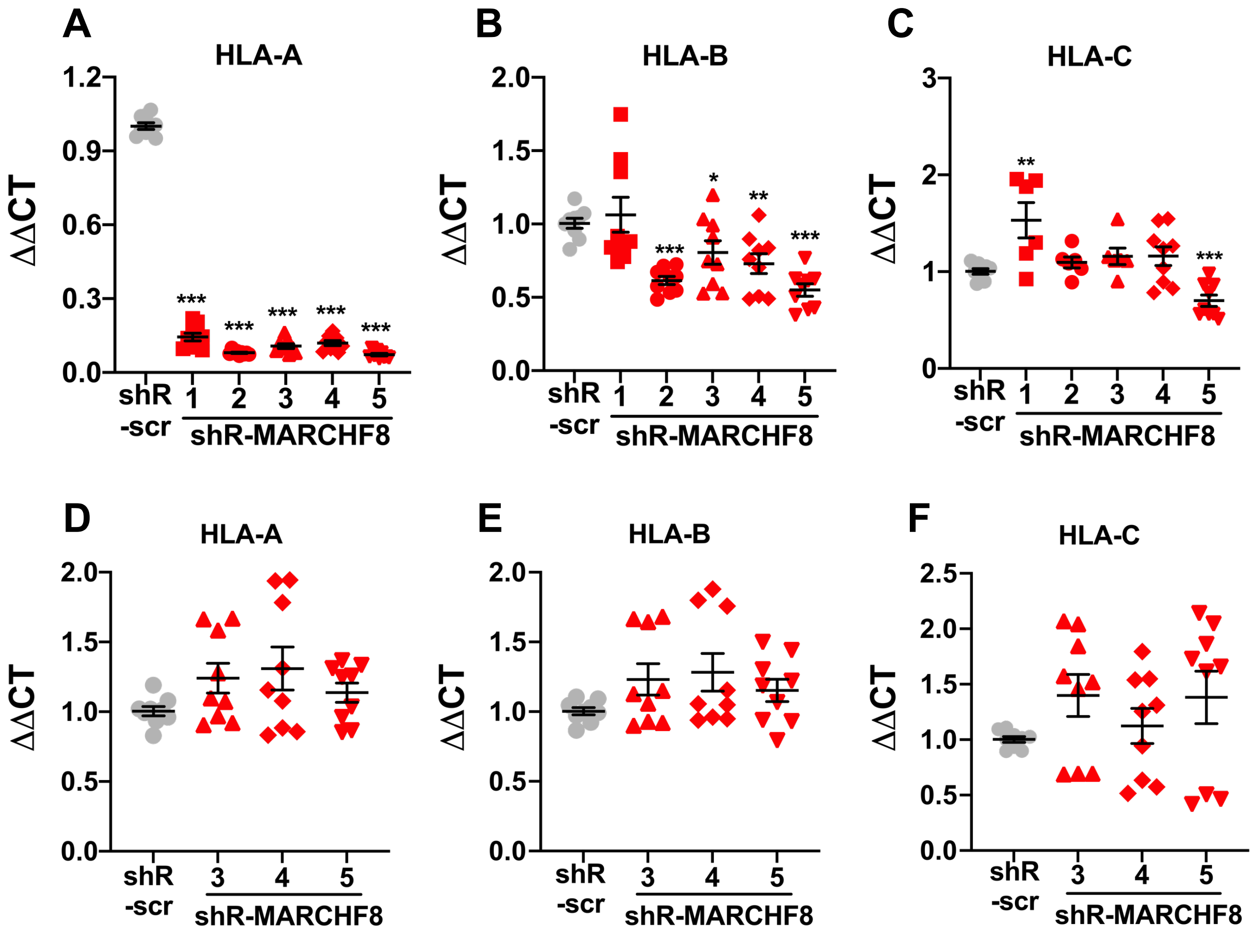


**Fig. S3. mRNA expression levels of HLA-A, HLA-B, and HLA-C in HPV+ HNC cells with MARCHF8 knockdown.** Two HPV+ HNC cell lines, SCC152 (**A** - **C**) and SCC2 (**D** - **F**), were transduced with five and three lentiviral shRNAs against MARCHF8 (shR-MARCHF8), respectively, or scrambled shRNA (shR-scr). The mRNA levels of HLA-A (**A** and **D**), HLA-B (**B** and **E**), and HLA-C (**C** and **F**) were assessed by RT-qPCR. The data shown are normalized by the GAPDH mRNA level as internal control. All experiments were repeated at least three times, and the data shown are means ± SD. *P* values were determined by Student's *t*-test. *$p < 0.05$, **$p < 0.01$, ***$p < 0.001$.

Khalil et al., Fig.S4

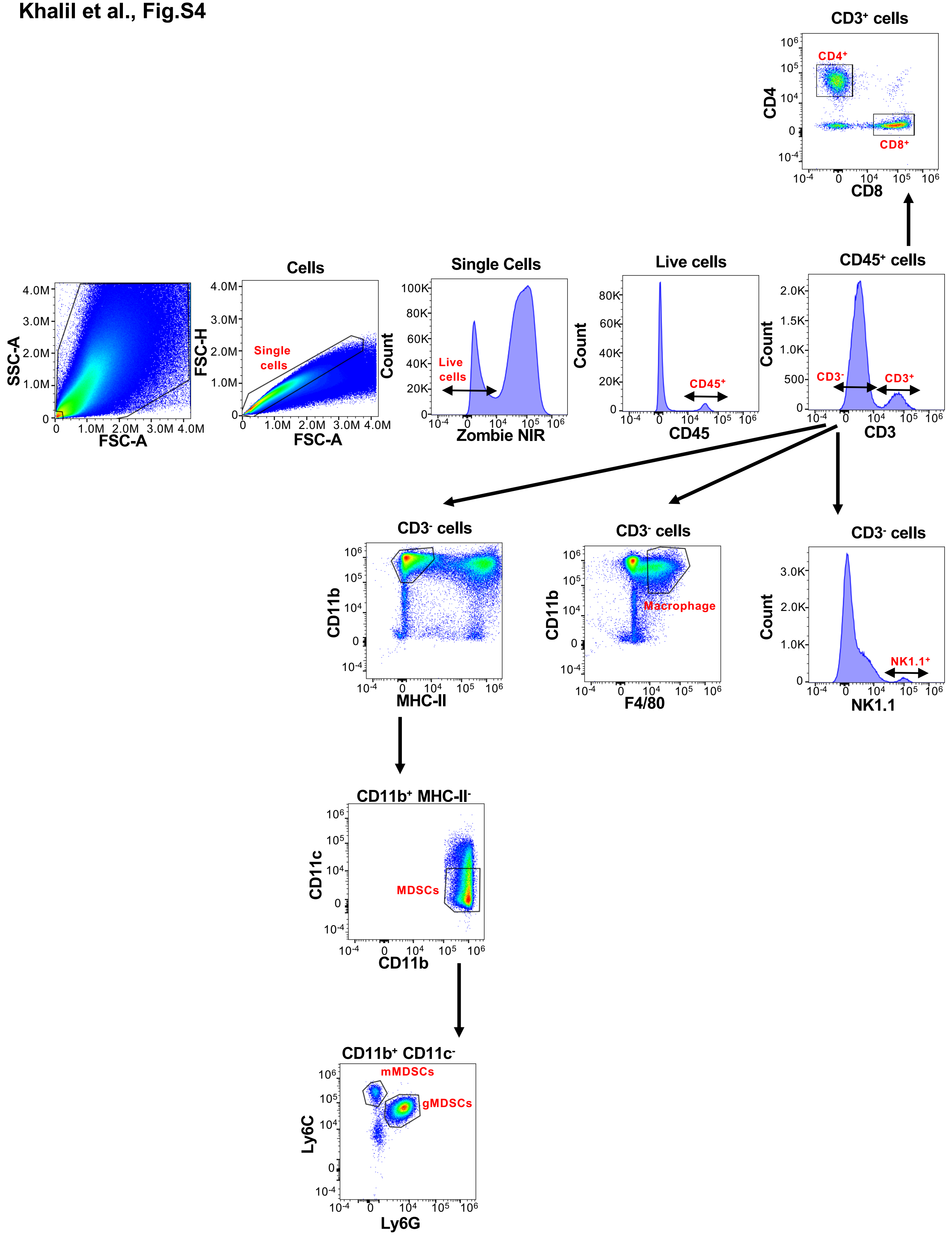

**Fig. S4. Representative gating strategy for flow cytometry analysis**. Single cells isolated from the tumor tissues were stained with an antibodies cocktail containing antibodies for CD45, CD3, CD4, CD8, CD11b, CD11c, MHC-II, F4/80, NK1.1, Ly6C, and Ly6G and the viability dye zombie NIR. The single cells were gated for live cells using zombie NIR viability dye and subsequently gated for CD45 expression as a marker for immune cells. $CD45^+$ immune cells were classified into $CD3^-$ and $CD3^+$ cells based on CD3 expression. The $CD3^+$ T cells were branched into $CD4^+$ and $CD8^+$ T cells, while $CD3^-$ cells contain macrophages ($F4/80^+$), NK cells ($NK1.1^+$), and MDSCs ($CD11b^+$ $MHCII^-$). MDSCs were divided into granulocytic (gMDSC, $Ly6G^+$) and monocytic (mMDSC, $Ly6C^+$).

Khalil et al., Fig. S5

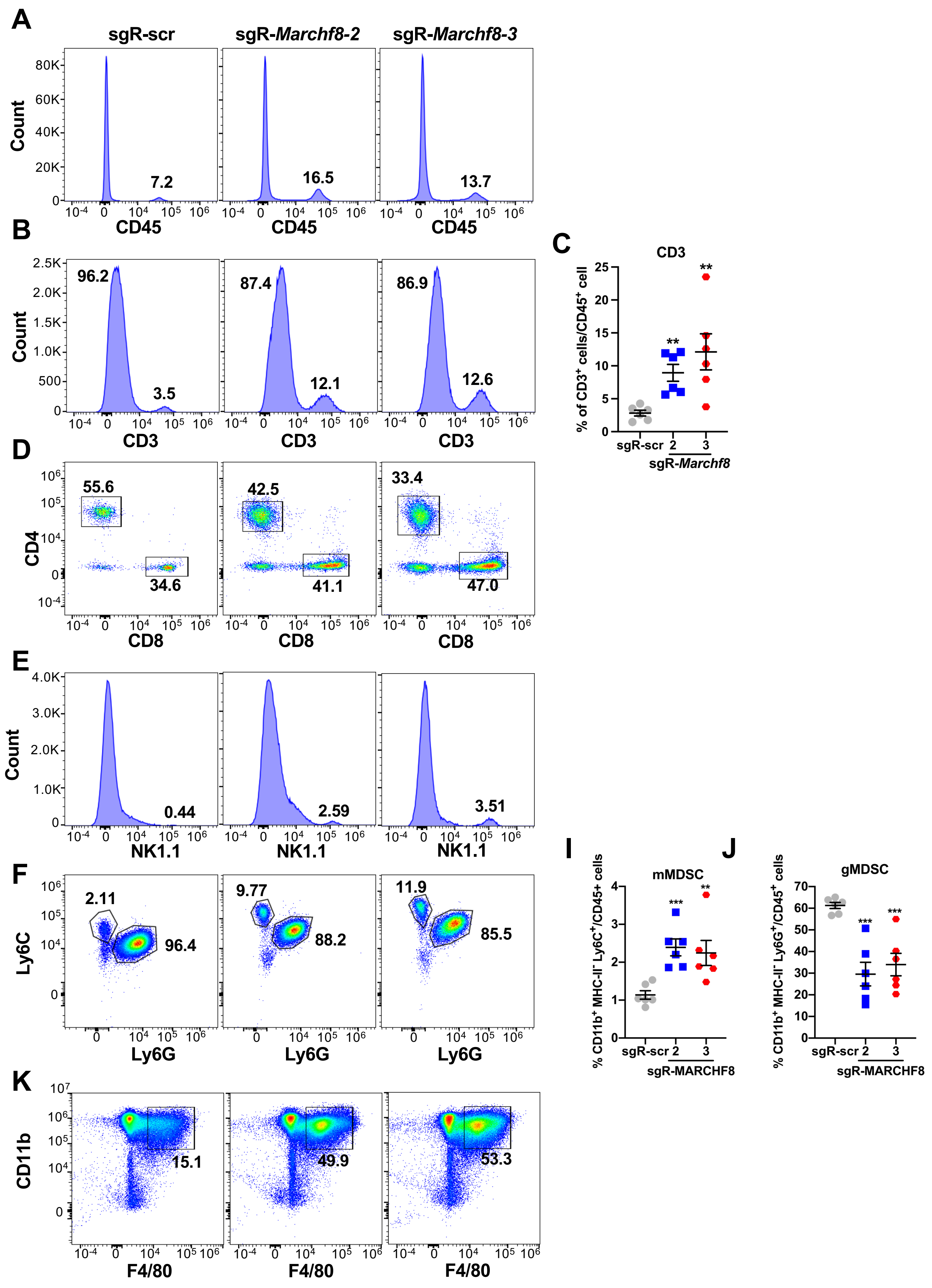

**Fig. S5. *Marchf8* knockout in HPV+ HNC cells increases NK cells and macrophages and decreases granulocytic MDSCs in the tumor tissues.** Tumors were isolated from C57BL/6J mice injected with mEERL/scr or mEERL/*Marchf8*$^{-/-}$ (sgR-Marchf8-2 and sgR-Marchf8-3) cells. Single cells isolated from the tumor tissues were stained with an antibody cocktail and analyzed by flow cytometry**.** Representative flow cytometry plots show CD45$^{+}$ immune cells (**A**), CD3$^{+}$ (**B** and **C**), CD4$^{+}$ and CD8$^{+}$ T cells (**D**), NK1.1$^{+}$ NK cells (**E**), mMDSCs (CD11b$^{+}$ MHC-II$^{-}$ Ly6C$^{+}$) (F and I), and gMDSCs (CD11b$^{+}$ MHC-II$^{-}$ Ly6G$^{+}$) (**F and J**), and macrophages (CD11b$^{+}$ F4/80$^{+}$) (**K**). MFI of three independent experiments is shown (**C, I, J**). All experiments were repeated at least three times, and the data shown are means ± SD. *P* values were determined by Student's *t*-test. $^{**}p < 0.01$, $^{***}p < 0.001$.

Khalil et al., Fig.S6

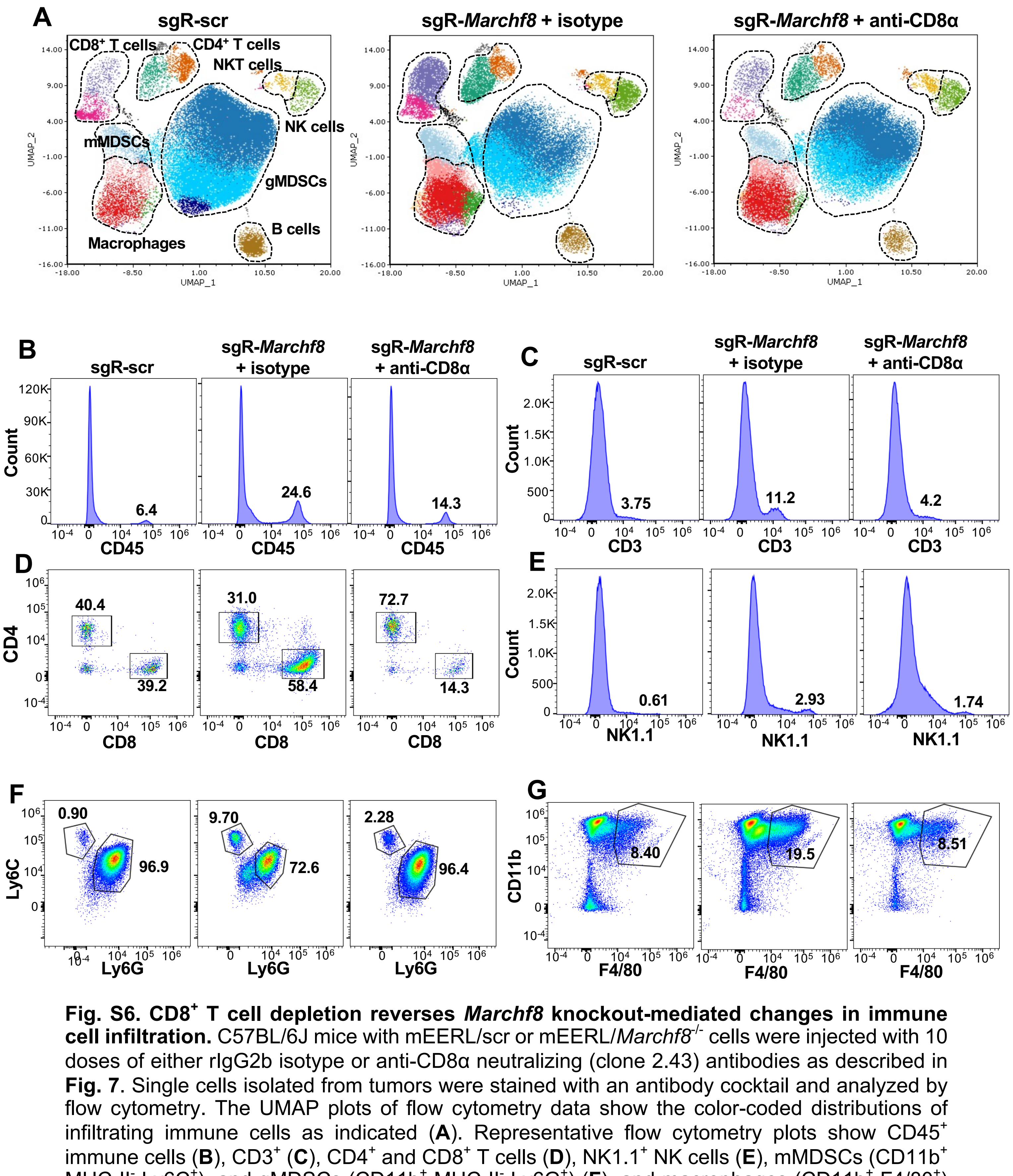


**Fig. S6. CD8$^{+}$ T cell depletion reverses *Marchf8* knockout-mediated changes in immune cell infiltration.** C57BL/6J mice with mEERL/scr or mEERL/*Marchf8*$^{-/-}$ cells were injected with 10 doses of either rIgG2b isotype or anti-CD8α neutralizing (clone 2.43) antibodies as described in **Fig. 7**. Single cells isolated from tumors were stained with an antibody cocktail and analyzed by flow cytometry. The UMAP plots of flow cytometry data show the color-coded distributions of infiltrating immune cells as indicated (**A**). Representative flow cytometry plots show CD45$^{+}$ immune cells (**B**), CD3$^{+}$ (**C**), CD4$^{+}$ and CD8$^{+}$ T cells (**D**), NK1.1$^{+}$ NK cells (**E**), mMDSCs (CD11b$^{+}$ MHC-II$^{-}$ Ly6C$^{+}$), and gMDSCs (CD11b$^{+}$ MHC-II$^{-}$ Ly6G$^{+}$) (**F**), and macrophages (CD11b$^{+}$ F4/80$^{+}$) (**G**). All experiments were repeated at least three times.

### Table S1. List of the antibodies

| Antibody | Specificity | Source | Catalog | RRID | Experiment |
|---|---|---|---|---|---|
| HPV16 E7 (clone ED17) | Virus | Santa Cruz | SC-6981 | AB_627745 | Western blot |
| MARCHF8 | Human/Mouse | Thermo Fisher | PA5-88893 | AB_2805201 | Western blot |
| MARCHF8 | Human/Mouse | Proteintech | 14119-1-AP | AB_2140168 | IP |
| HLA class I ABC | Human | Proteintech | 15240-1-AP | AB_1557426 | IP-WB |
| ubiquitin | Human/Mouse | Proteintech | 10201-2-AP | AB_671515 | IP |
| FITC-HLA-A, B, C | Human | BioLegend | 311404 | AB_314872 | Flow cytometry |
| PE-H-2Ld/H-2Db | Mouse | BioLegend | 114507 | AB_313588 | Flow cytometry |
| Anti-mouse CD8α (clone 2.43) | Mouse | BioXCell | BE0061 | AB_1125541 | CD8 T cell depletion |
| Anti-mouse PD-1 (clone RMP1-14) | Mouse | BioXCell | BE0146 | AB_10949053 | PD-1 blockade |
| Rat IgG2a isotype control | Mouse | BioXCell | BE0089 | AB_1107769 | Isotype control |
| Rat IgG2b isotype control | Mouse | BioXCell | BE0090 | AB_1107780 | Isotype control |
| Brilliant Violet 510-Ly6C | Mouse | BioLegend | 128033 | AB_2562351 | Flow cytometry panel |
| PerCP-Ly6G | Mouse | BioLegend | 127653 | AB_2616999 | Flow cytometry panel |
| PE/Dazzle 594 I-A/I-E | Mouse | BioLegend | 107647 | AB_2565979 | Flow cytometry panel |
| APC/Fire 750-CD8a | Mouse | BioLegend | 100765 | AB_2572113 | Flow cytometry panel |
| Brilliant Violet 605-F4/80 | Mouse | BioLegend | 123133 | AB_2562305 | Flow cytometry panel |
| Brilliant Ultraviolet 395-CD45 | Mouse | Thermo Fisher | 363-0451-80 | AB_2925263 | Flow cytometry panel |
| Super Bright 780-CD11b | Mouse | Thermo Fisher | 78-0112-80 | AB_2722925 | Flow cytometry panel |
| APC/Fire 810-CD3 | Mouse | BioLegend | 100268 | AB_2876392 | Flow cytometry panel |
| PE/Cy7-NK1.1 | Mouse | BioLegend | 108714 | AB_389364 | Flow cytometry panel |
| PE-CD11c | Mouse | BioLegend | 117308 | AB_313777 | Flow cytometry panel |
| FITC-CD4 | Mouse | BioLegend | 100510 | AB_312713 | Flow cytometry panel |
| Spark Blue 550-CD19 | Mouse | BioLegend | 115565 | AB_2819827 | Flow cytometry panel |
| PE/Fire 810-Tim3 | Mouse | BioLegend | 119745 | AB_2922462 | Flow cytometry panel |
| APC-LAG3 | Mouse | BioLegend | 125209 | AB_10639935 | Flow cytometry panel |
| PerCP-eFluor 710-PD-1 | Mouse | Thermo Fisher | 46-9981-80 | AB_11149347 | Flow cytometry panel |

RRID, Research Resource Identifiers

**Table S2. List of the oligonucleotides**

| Name | Sequence | Experiment |
|---|---|---|
| Human GAPDH FWD | 5`-GGAGCGAGATCCCTCCAAAAT-3` | RT-qPCR |
| Human GAPDH Rev | 5`-GGCTGTTGTCATACTTCTCATGG-3` | RT-qPCR |
| Human HLA-A FWD | 5`-CTTGTAAAGTGTGAGACAGC-3` | RT-qPCR |
| Human HLA-A Rev | 5`-CTTCAAGTCACAAAGGGAAG-3` | RT-qPCR |
| Human HLA-B FWD | 5`-ATGTGTAGGAGGAAGAGTTC-3` | RT-qPCR |
| Human HLA-B Rev | 5`-GAAGAAATCCTGCATCTCAG-3` | RT-qPCR |
| Human HLA-C FWD | 5`-CATCACTTGTAAAGCCTGAG-3` | RT-qPCR |
| Human HLA-C Rev | 5`-CTCTTGAAGTCACAAAGGAG-3` | RT-qPCR |
| 3X HA tag FWD | 5`-TCCGGAACTAGTATGATCTTTTACCCATAC-3` | Cloning |
| Human MARCHF8 Rev | 5`-TTACTAACCGGTTCAGACGTGAATGAT-3` | Cloning |
| Human MARCHF8 1-220 Rev | 5`-TTACTAACCGGTTCATTTACACTGAACATACATAAA-3` | Cloning |
| Human MARCHF8 1-250 Rev | 5`-TTACTAACCGGTTCAAAAAATATTCTTTTTGCTTGT-3` | Cloning |
| Human MARCHF8 1-270 Rev | 5`-TTACTAACCGGTTCAATGACAGATTCCATATCCATG-3` | Cloning |
| Human MARCHF8 ∆TM1 FWD | 5`-GGAAGATCATGTGCTCAGTGACAGTGCTCATTGACCGT ACTGCTG-3` | Cloning |
| Human MARCHF8 ∆TM1 Rev | 5`-CAGCAGTACGGTCAATGAGCACTGTCACTGAGCACATGATCTTCC-3` | Cloning |
| Human MARCHF8 ∆TM2 FWD | 5`-TAGAATGGCCCTTTTGGACTAAAATGTATGTTCAGTGTAAAGTGC-3` | Cloning |
| Human MARCHF8 ∆TM2 Rev | 5`-GCACTTTACACTGAACATACATTTTAGTCCAAAAGGGCCATTCTA-3` | Cloning |
| Human MARCHF8 ∆DIRT FWD | 5`-CATGGAGACCAAGCTGAAGCCAAGGAAGATCATGTGCT CAGTGAC-3` | Cloning |
| Human MARCHF8 ∆DIRT Rev | 5`-GTCACTGAGCACATGATCTTCCTTGGCTTCAGCTTGGTCTCCATG-3` | Cloning |

FWD, forward primer; Rev, reverse primer

### Table S3. List of the shRNAs

| Name | Sigma-Aldrich TRC Clone ID | Sequence |
|---|---|---|
| Human MARCHF8 shRNA1 | TRCN0000073233 | 5`-CTTGAGCTGAATGAGAGAATA-3` |
| Human MARCHF8 shRNA2 | TRCN0000073234 | 5`-CCACTAACAGAGCCCAACTTT-3` |
| Human MARCHF8 shRNA3 | TRCN0000073235 | 5`-CAGTGTAAAGTGTATGTGCAA-3` |
| Human MARCHF8 shRNA4 | TRCN0000073236 | 5`-CTGGTCCTTGTATGTGCTCAT-3` |
| Human MARCHF8 shRNA5 | TRCN0000073237 | 5`-CCTCCTTCTCTCGCACTTCTA-3` |

### Table S4. List of the sgRNAs

| Name | Sequence |
|---|---|
| Mouse MARCHF8 sgRNA1 FWD | 5`-CACCGAGGTGAGTATATGGGCCGTGAGG-3` |
| Mouse MARCHF8 sgRNA1 Rev | 5`-AAACCCTCACGGCCCATATACTCACCT-3` |
| Mouse MARCHF8 sgRNA2 FWD | 5`-CACCGTATTAACGTCTGACCATGTGAGG-3` |
| Mouse MARCHF8 sgRNA2 Rev | 5`-AAACCCTCACATGGTCAGACGTTAATA-3` |
| Mouse MARCHF8 sgRNA3 FWD | 5`-CACCGACTACCAGCTTCGTCCAGAAAGG-3` |
| Mouse MARCHF8 sgRNA3 Rev | 5`-AAACCCTTTCTGGACGAAGCTGGTAGT-3` |

FWD, forward oligo; Rev, reverse oligo
